# Isogenic cortical organoids enable precision targeting of APP variant-specific pathways in Alzheimer’s disease

**DOI:** 10.64898/2026.02.19.706886

**Authors:** T Grass, IM Cosacak, A Ordureau, FD Price, S Kavali, A Caldarelli, MN Qiao, B Vardarajan, Joao A. Paulo, Michele Marass, LL Rubin, C Kizil, N Rodriguez-Muela

**Affiliations:** German Center for Neurodegenerative Diseases e.V. (DZNE), Dresden, Germany; Technische Universität Dresden, Center for Regenerative Therapies Dresden, Dresden, Germany; Cell Biology Program, Sloan Kettering Institute, Memorial Sloan Kettering Cancer Center, New York, NY, USA; Department of Stem Cell and Regenerative Biology, Harvard University, Cambridge, MA, USA; The Taub Institute for Research on Alzheimer’s Disease and the Aging Brain, Columbia University Irving Medical Center, Columbia University, New York, NY, USA; Department of Neurology, Columbia University Irving Medical Center, Columbia University, New York, NY, USA; Department of Cell Biology, Blavatnik Institute, Harvard Medical School, Boston, MA, USA; Center for Systems Biology Dresden, Dresden, Germany; Max Planck Institute for Molecular Cell Biology and Genetics, Dresden, Germany; Stanley Center for Psychiatric Research, Broad Institute of MIT and Harvard, Cambridge, MA, USA; Harvard Stem Cell Institute, Cambridge, MA, USA

## Abstract

Alzheimer’s disease (AD) lacks disease-modifying therapies, in part due to the limitations of existing disease models, which have struggled to capture the early pathogenic events leading to neuronal degeneration. Unfortunately, recent therapies targeting hallmarks of AD have proven inefficient in humans, and it is thus necessary to identify alternative targets. Here, by generating an isogenic panel of hiPSC-derived cortical organoids carrying familial AD-associated *APP* variants or the protective A673T variant, we identified distinct, actionable pathogenic pathways specific to each variant. Proteomic analyses revealed variant-specific molecular disruptions: A673V organoids show impairments in proteostasis and cholesterol metabolism, whereas KM670/671NL organoids exhibit mitochondrial bioenergetic defects. These signatures overlapped with dysregulated proteins in post-mortem AD brains, demonstrating the reliability of our in vitro model. Importantly, targeted interventions restored neuronal survival in a variant-specific manner: overexpression of the master regulator of lysosomal biogenesis, TFEB, rescued A673V neurons, while ferroptosis inhibition selectively protected KM670/671NL neurons. Overall, our results indicate that differential treatments can be tailored based on distinct genetic backgrounds, supporting the development of precision medicine approaches in AD.

## Introduction

Alzheimer’s disease (AD) is the leading cause of dementia - which is ranked as the 7th leading cause of death globally - and the most prevalent neurodegenerative disease (ND). Currently, over 55 million people are affected worldwide ^1,2^, and this number is projected to triple by 2050 ^3^. Despite decades of research and numerous therapeutic trials targeting amyloid-β (Aβ) and tau ^4,5^, no treatment has yet prevented or reversed disease progression, and existing therapies remain largely symptomatic ^6^. Recent advances with anti-Aβ monoclonal antibodies have shown that reducing amyloid burden can modestly slow cognitive decline ^5,7,8^, supporting Aβ dysregulation as a key pathogenic mechanism^4^. However, these findings also emphasize the need to intervene before significant neurodegeneration occurs. Because Aβ is produced through sequential cleavage of the amyloid precursor protein (APP)^9^, the partial efficacy of these therapies reinforces the centrality of APP processing in AD pathogenesis and highlights the need to understand how early disruptions in APP metabolism arise. Such insight may help to identify the initial molecular events that precede overt neurodegeneration, particularly as AD-related brain changes are thought to begin 20 years or more before clinical symptom onset ^10^.

While extracellular Aβ plaques and intracellular neurofibrillary tangles composed of hyperphosphorylated tau (MAPT) are well-established hallmarks of AD ^11,12^, the initiating molecular cascades and mechanisms underlying selective neuronal vulnerability remain incompletely understood. Post-mortem human tissue provides only a snapshot of late-stage pathology, and although animal and overexpression models have been indispensable in elucidating aspects of AD biology, they only partially recapitulate human-specific disease mechanisms and have shown limited translational value ^13,14^. This has created an urgent need for human-relevant model systems capable of capturing early, causal molecular changes.

Human induced pluripotent stem cell (hiPSC) models provide a powerful platform to investigate early, human-specific pathogenic processes in a genetically controlled context ^15,16^. The use of isogenic hiPSC lines further enhances causal inference by eliminating inter-individual genetic variability, enabling precise comparison of disease-associated alleles. Growing evidence from hiPSC-derived neurons and organoids has shown that AD-linked mutations can disrupt early neuronal maturation, proteostasis, and mitochondrial function well before the appearance of extracellular Aβ plaques or intracellular tau tangles ^17,18^. These findings challenge the classical view of Aβ accumulation as a purely late-stage event and suggest that early intracellular Aβ and associated stress responses may prime neurons for degeneration later in life.

Motivated by this concept, we sought to identify early transcriptional and translational signatures associated with distinct *APP* variants that may foreshadow later AD pathology. Using CRISPR–Cas9 genome editing, we generated an isogenic panel of hiPSC lines carrying *APP* variants linked to early-onset AD, including the Swedish double mutation (K670M/N671L), the A673V mutation as well as the protective Icelandic A673T variant - associated with reduced susceptibility to AD and preserved cognitive function during aging-, alongside wild-type and *APP* knockout controls ^19–23^. These mutations differentially affect APP processing ^24,25^. Specifically, A673V -the only known recessive *APP* mutation- lies within the Aβ sequence (residue 2), increasing Aβ aggregation kinetics and fibrillogenic potential ^22,26^. In contrast, the dominant Swedish mutation is located just outside the Aβ region at the β-secretase cleavage site, increasing BACE1 accessibility and elevating production of wild-type Aβ ^20,27,28^. Despite acting through distinct mechanisms, both mutations shift β-secretase cleavage from the physiological Glu11 (C89) site (β’-site) to the amyloidogenic Asp1 (C99) site (β-site), increasing the pool of C99 fragments 29,30. Notably, downstream processing also diverges: C99 derived from A673V is preferentially routed to proteasomal degradation, while Swedish APP results in higher steady-state Aβ levels due to enhanced amyloidogenic flux ^29^. These complementary biochemical profiles make A673V and SWE an ideal pair for probing how qualitatively distinct disruptions in APP processing shape early aspects of AD biology. On the contrary, the Icelandic variant increases the turnover rate of APP by α-secretase, known as amyloidolytic APP processing, as cleavage occurs within the Aβ sequence precluding its generation. Additionally, it promotes cleavage of BACE-1 at β’-site^23,31^. Using this genetically controlled system, we then mapped early proteomic, transcriptional, and phenotypic changes to determine whether these variants activate shared molecular programs or instead reveal mutation-specific vulnerabilities.

We found that *APP* variants elicit distinct dysregulation of pathways essential to neuronal health, particularly those related to proteostasis, lysosomal function, and mitochondrial bioenergetics. Our model recapitulates key features of AD, including intracellular and extracellular Aβ accumulation and reduced neuronal survival, while also revealing mutation-specific impairments in protein homeostasis and mitochondrial metabolism. Several dysregulated proteins overlapped with those identified in post-mortem AD brains ^32,33^, highlighting new candidate pathways with translational relevance. Together, these findings demonstrate that distinct *APP* variants drive both shared and unique early molecular changes that link initial stress responses to later neurodegenerative cascades. This work provides mechanistic insight into the earliest stages of human AD and supports the use of isogenic hiPSC-derived organoid models for studying variant-specific vulnerabilities and informing early-stage, precision therapeutic strategies.

## Results

### hiPSC-derived cortical neurons carrying familial AD mutations show reduced survival

AD is marked by the selective death of certain neuronal subtypes, particularly excitatory neurons in the entorhinal cortex expressing RORB and reelin and somatostatin (SST) - expressing interneurons in the association cortex -, and pyramidal neurons in the CA1 region of the hippocampus ^34–38^. To investigate neuronal vulnerability in fAD, we used an isogenic hiPSC model that we newly developed carrying different *APP* variants introduced via CRISPR/Cas9-mediated genome editing into a healthy (wild type, WT) hiPSC line (BJSipsD), using a two-vector-based targeting-approach ^39^ (Figure S1A-B). We established two hiPSC lines harboring pathogenic fAD mutations: A673V mutation ^22^ (“AV” for short) and the Swedish KM670/671NL double mutation ^20,21^ (“SWE” for short), both associated with early-onset AD. In addition, we generated a line carrying the only reported protective variant in *APP*, the Icelandic A673T variant ^19^ (“AT” for short), which reduces the risk of AD and age-related cognitive decline, as well as an *APP* knock-out (*APP* KO) line (Figure S1C). Successfully edited clones were identified by PCR-based genotyping and Sanger sequencing (Figure S1D). All lines maintained normal karyotype post-editing, showed no detectable off-target edits at the top 5 candidate sites (data not shown) and retained pluripotency (Figure S1E-G). Western blot analysis confirmed APP protein expression in all edited lines except for the KO (Figure S1H-I). Interestingly, APP levels were significantly reduced in the AT and AV lines compared to WT, with a downward trend also observed in the SWE line, possibly reflecting increased α-and β-secretase cleavage activity in the presence of the AT, AV, and SWE variants, respectively.

Once our isogenic lines were validated, we first explored whether the fAD-linked *APP* mutations impaired cortical neuron (CxN) survival in vitro. Following a well-established protocol ^40^ (Figure 1A), we generated cortical spheres, dissociated them at day 45 of differentiation, and plated the resulting CxNs. Neuronal survival was then live-tracked over time using calcein red dye and an automated high-content imaging microscope. This system enabled standardized, unbiased comparison of variant-dependent differences in neuronal resilience. Automated, unbiased quantification of live fluorescence in hundreds of thousands of neurons revealed that healthy (WT) CxNs survived significantly better than those carrying the fAD-AV mutation or lacking APP, while the CxNs carrying the protective AT variant showed significantly greater survival than AV, SWE and KO neurons (Figure 1B-C). We further validated that these cultures consisted predominantly of neurons, as confirmed by positive staining for the pan-neuronal marker MAP2, and of deep-layer excitatory neurons, including CTIP2-expressing subcerebral projection neurons, a population known to be selectively vulnerable in early AD ^34^ (Figure 1D-H). In line with our calcein red live-tracking data, we observed that significantly more WT CxNs survived over a two-week period compared to both the fAD and KO CxNs. Interestingly, this analysis also revealed that AT CxNs, carrying the protective variant, were notably more resilient than WT CxNs under the same culture conditions (Figure 1D-F). To rule out that a different neuronal composition of the spheres carrying the distinct *APP* mutations was responsible for the diminished CxN survival in the fAD or the *APP* KO cultures, we quantified the percentage of MAP2^+^ and CTIP2^+^ CxNs shortly after cortical sphere dissociation and plating. We did not detect significant differences across the lines, which were composed of ∼95% MAP2^+^ neurons, of which 30-40% were CTIP2^+^ (Figure 1G-H). Given that dissociation and plating at the end of the differentiation protocol and subsequent plating of the dissociated neurons can be a harsh procedure that may disproportionately affect vulnerable cells, we further assessed whether fAD-associated mutations or *APP* loss impaired the ability of neural progenitors to differentiate into CxNs. To this end, we sought to quantify CTIP2^+^ neurons right after the dissociation of the cortical spheres. To do so, we generated reporter lines by C-terminally tagging the *CTIP2* locus with a P2A signal followed by the fluorescent green protein mNeonGreen (Figure S2A-B). Successfully targeted clones were validated by PCR and Sanger sequencing, and the specificity of the reporter was confirmed by co-localization of the endogenous mNeonGreen signal with anti-CTIP2 antibody immunostaining in dissociated cortical cultures (Figure S2C-F). Once validated, cortical spheres from the isogenic fAD CTIP2 reporter lines were generated and dissociated after 45 days as above, and the dissociated neurons analyzed by FACS. No significant differences in the percentage of CTIP2:mNeonGreen^+^ neurons were observed across genotypes (Figure S2 G-H), confirming that neither the pathogenic *APP* mutations nor *APP* knock out impair the differentiation potential of neuronal progenitors into CTIP2^+^ CxNs. Instead, these findings suggest that CxNs carrying the protective *APP* variant are more resilient to intrinsic culture-related stress, whereas those harboring fAD-associated mutations are more vulnerable. Overall, our results indicate that, in our isogenic system, neurons with deleterious APP variants exhibit vulnerability to cell death, whereas neurons carrying a protective APP variant show resilience to cell death.

**Figure 1.**
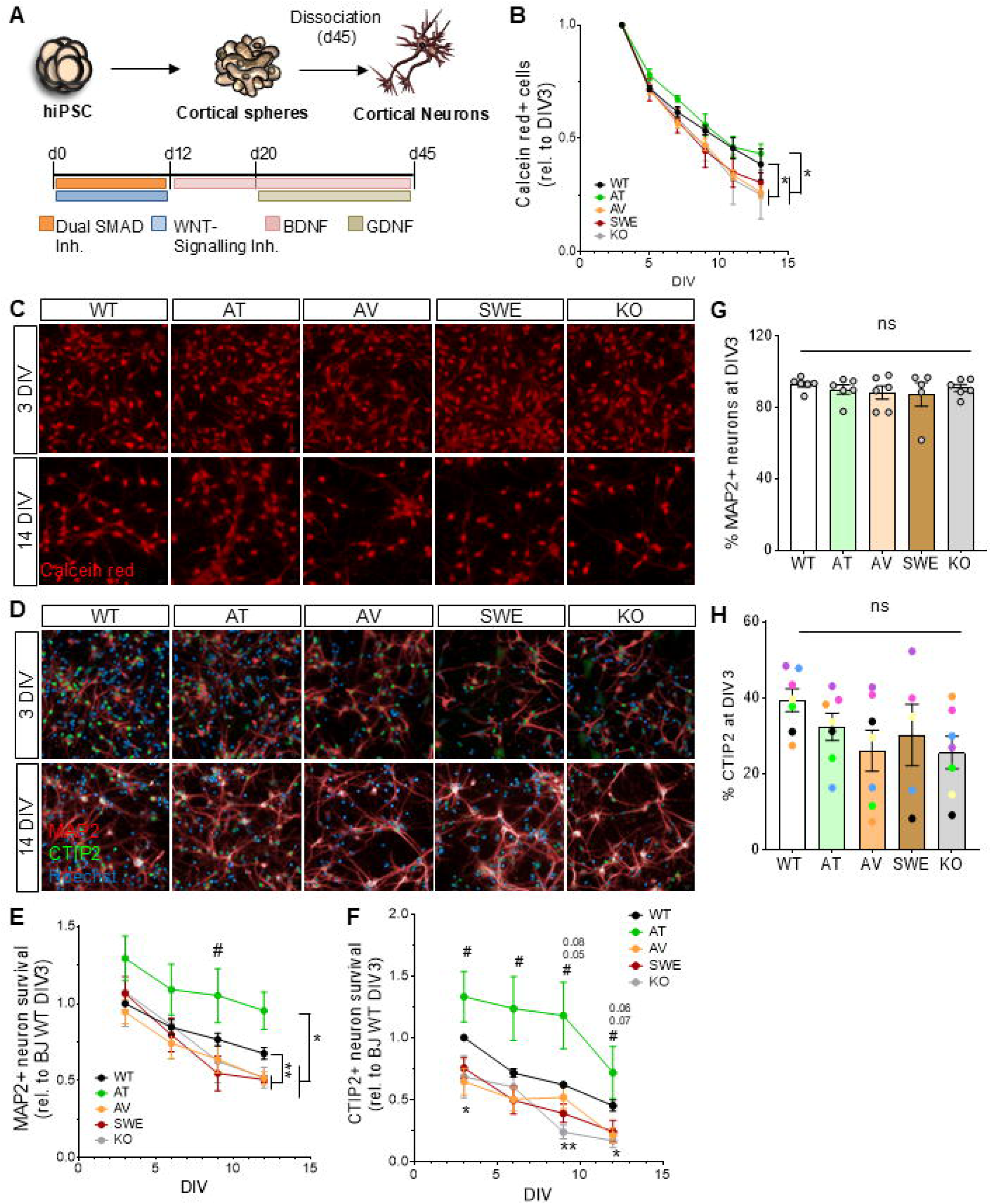
hiPSC-derived cortical neurons carrying familial AD mutations show reduced survival. **(A)** Schematic representation of the protocol followed to differentiate the fAD hiPSCs lines into CxNs. (**B**) Quantification of the number of cortical neurons (CxNs) labeled with the live dye Calcein red at the indicated time-points after the cortical spheres are dissociated and plated. Shown values are relative to day 3 for each line. (**C**) Representative images showing CxNs from all 6 hiPSC lines, 3 and 14 days after plating and live stained with Calcein red. Scale bar, 10 µm. (**D**) Representative images showing CxNs 3 and 14 days after plated, fixed and immunostained against MAP2 (red) and CTIP2 (green). Nuclei are labeled with Hoechst (blue). Scale bar, 10 µm. (**E**) Quantification of the number of CxNs immunostained against the pan-neuronal marker MAP2 at the indicated time-points after plating and fixation. Shown values are relative to day 3 healthy control BJ WT line. (**F**) Quantification of the number of CxNs immunostained against the CxN marker CTIP2 at the indicated time-points after plating and fixation. Shown values are relative to day 3 healthy control BJ WT line. Quantification of the percentage of MAP2^+^ (**G**) and CTIP2^+^ (**H**) CxNs out of all cells in the cultures at an early time-point (3 days after the cortical spheres are dissociated and plated). (One-way ANOVA followed by Bonferroni’s Multiple Comparison Test).

### Familial AD cortical neuron cultures show increased levels of Aβ40 and Aβ42 peptides

To confirm that our isogenic AD hiPSC lines recapitulate variant-dependent alterations in APP cleavage, we examined whether the introduced *APP* variants influenced the cleavage efficiency of the α- and β-secretases, thereby decreasing Aβ40-42 production in the AT line or increasing it in the AV and SWE lines, as expected. ELISA measurements of Aβ40 in the culture medium showed significantly lower levels in AT hiPSCs compared to WT, and nearly threefold higher levels in SWE hiPSCs. No significant differences were observed in the AV line (Figure 2A). The levels of the less abundant Aβ42 peptide were comparable between the protective AT and the control lines but were elevated by approximately 2 to 2.5-fold in the AV and SWE lines, respectively (Figure 2B). As expected, no Aβ peptides were detected in the KO line, which served as background control in these studies. These results indicate that *APP* fAD mutations in pluripotent stem cells result in aberrantly elevated Aβ42 production, and in the case of the SWE mutation, also Aβ40, which could lead to the dysregulation of cellular homeostasis. We next investigated whether similar changes in Aβ production were observed in CxNs. As expected, the amounts of both Aβ peptides in the cultures were approximately 3x and 5x higher in the fAD-AV and fAD-SWE CxNs compared to WT cells, respectively, while reduced levels were detected in the AT CxNs (Figure 2C-D). Again, Aβ40-42 levels were undetectable in *APP* KO CxNs (Figure 2C-D). Altogether, these results confirm that the AD-linked mutations introduced in the *APP* locus led to the expected altered production of the prone-to-aggregate Aβ peptides and validate our human in vitro AD model.

**Figure 2.**
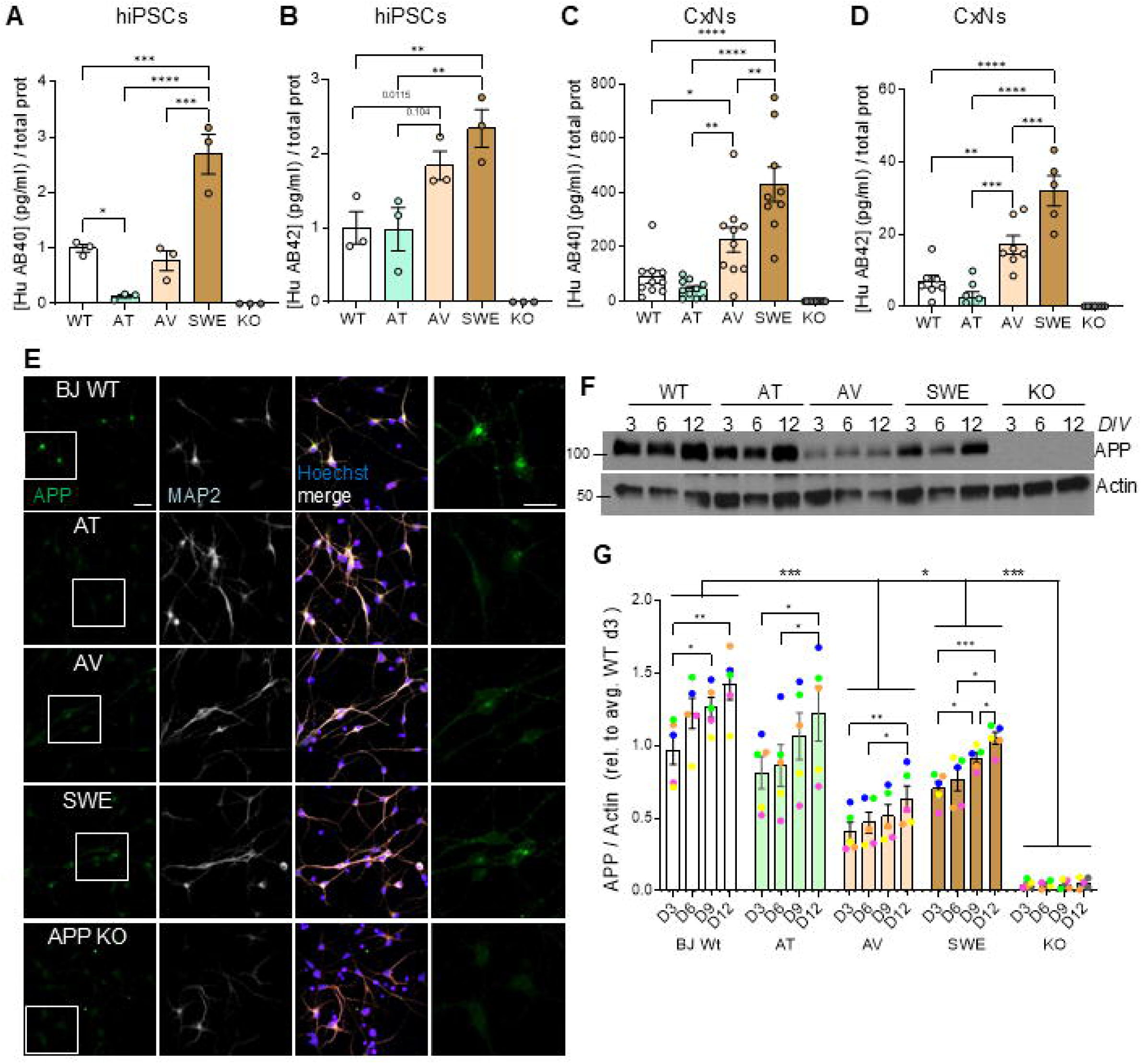
Familial AD lines show increased levels of Aβ40 and Aβ42 peptides. ELISA quantification of Aβ40 (**A**) and Aβ42 (**B**) *APP*-derived peptides in the culture media from the isogenic fAD hiPSCs. ELISA quantification of Aβ40 (**C**) and Aβ42 (**D**) in the culture media from CxNs plated from 12 days after day45 cortical sphere dissociation (One-way ANOVA followed by Bonferroni’s Multiple Comparison Test). (**E**) Representative confocal images of CxNs dissociated from day 45 spheres and kept in culture for 6 more days. Neurons were fixed and immunostained for *APP* (green) and MAP2 (white). Nuclei are stained with Hoechst (blue). A magnification of the squared region is shown on the far-right column. Scale bars, 20 µm. (**F**) Representative western blot from isogenic hiPSC-derived CxN cultures 3, 6 or 12 days after day-45 cortical sphere dissociation and plating showing *APP* protein levels and quantification (**G**). (Two-way ANOVA followed by Tukey’s Multiple Comparison Test).

To determine whether increased basal production of Aβ peptides in the fAD lines compared to the WT and AT correlates with reduced levels of APP protein in CxN cultures, we performed immunostaining for *APP*, MAP2, and CTIP2 on neurons derived from dissociated cortical spheres at multiple time points after plating (Figure 2E). Using automated, unbiased high-content imaging and image analysis^41^, the cytoplasm of the MAP2^+^ and CTIP2^+^ neurons was identified, and the intensity of APP fluorescence was quantified in this cell region. We observed that APP levels were highest in WT neurons, reduced in the other three AD lines, and absent in the APP KO cells (Figure 2E). As mRNA levels remained comparable across all lines (data not shown), these findings suggest that *APP* turnover or proteolytic cleavage – via α-secretase (ADAM10) in the AT line, and via β-secretase (BACE1) in the AV and SWE lines – is accelerated, leading to faster processing and correspondingly lower detectable *APP* levels in the plasma membrane and cytoplasm. These observations are consistent with our western blot results from undifferentiated hiPSCs (Figure S1H-I). Notably, APP levels in CxNs seem to increase over time in culture, regardless of the genotype, in agreement with previously published studies ^42^ (Figure 2F-G). These results highlight how our isogenic platform reproduces the pathogenic phenotypes associated with each APP variant observed in AD patients across scales.

### Isogenic familial AD cortical neurons are amenable to treatments

To determine whether our isogenic hiPSC-derived AD model is amenable to pharmacological modulation of Aβ production – and thus suitable for drug screening applications to identify potential disease modifiers – we tested the effects of β- and γ-secretase inhibitors, which block sequential APP cleavage. Day 45 cortical spheres were dissociated, and the resulting CxNs were plated and treated with either β-secretase inhibitor (β-secretase inhibitor IV) or the γ-secretase inhibitor DAPT every 3 days for a total of 12 days (Figure 3A). The quantification of Aβ40 levels in the cultured media by ELISA showed a significant reduction for both treatments compared to DMSO-treated cells across all lines (Figure 3B). Furthermore, β- and γ-secretase inhibitors significantly reduced Aβ42 production in the fAD AV and SWE CxNs to levels comparable to the ones observed for the WT and AT neurons (Figure 3B-C). Furthermore, β-and γ-secretase inhibitors significantly reduced Aβ42 production in the fAD AV and SWE CxNs to levels like those in WT and AT neurons (Figure 3B-C). However, the reduction appeared less pronounced than for Aβ40, likely because Aβ40 is the predominant γ-secretase product, produced at higher levels and more soluble, leading to a rapid loss upon inhibition ^43,44^. To confirm that secretase inhibition altered APP processing, we quantified total APP levels after the treatments by western blot. In day 45 CxNs treated with the β-secretase inhibitor, full-length APP significantly accumulated across all genotypes (Figure 3D-E), consistent with reduced cleavage by BACE1. Additionally, as expected, the γ-secretase inhibitors DAPT and CompE did not significantly change *APP* levels since γ-secretase acts downstream of β- or α-secretase cleavage (Figure 3A, D-E). We next assessed phosphorylation of tau at Ser202/Thr205 (AT8 epitope) and Ser396/404 (PHF1 epitope) (Figure 3D). No significant differences in phospho-tau levels were observed across AV or SWE CxNs or upon treatment, likely reflecting the early developmental stage of these cultures prior to the onset of robust tau hyperphosphorylation typically seen in more advanced models or post-mortem AD tissue ^45^. Next, we investigated whether the decrease in Aβ40-42 production following the secretase inhibition would impact the survival of CxNs carrying fAD *APP* mutations. Importantly, treatment with the BACE-1 inhibitor significantly increased the number of CTIP2^+^ CxNs in both AV and SWE cultures (Figure 3F). No pro-survival effect was observed in the KO line, consistent with the absence of APP-derived toxic species in these neurons. Notably, γ-secretase inhibition (CompE) led to a reduction in CTIP2+ neurons in WT and AT cultures (Figure 3F), likely reflecting increment of β-C-terminal fragments (β-CTFs), previously reported to be neurotoxic ^46^. In conclusion, these results demonstrate that our isogenic fAD cortical neuron model is responsive to pharmacological modulation of APP processing and Aβ production. The ability to rescue neuronal survival in mutation carriers via β-secretase inhibition highlights the utility of this platform for studying disease mechanisms and screening for candidate therapeutic interventions in a genetically controlled human context.

**Figure 3.**
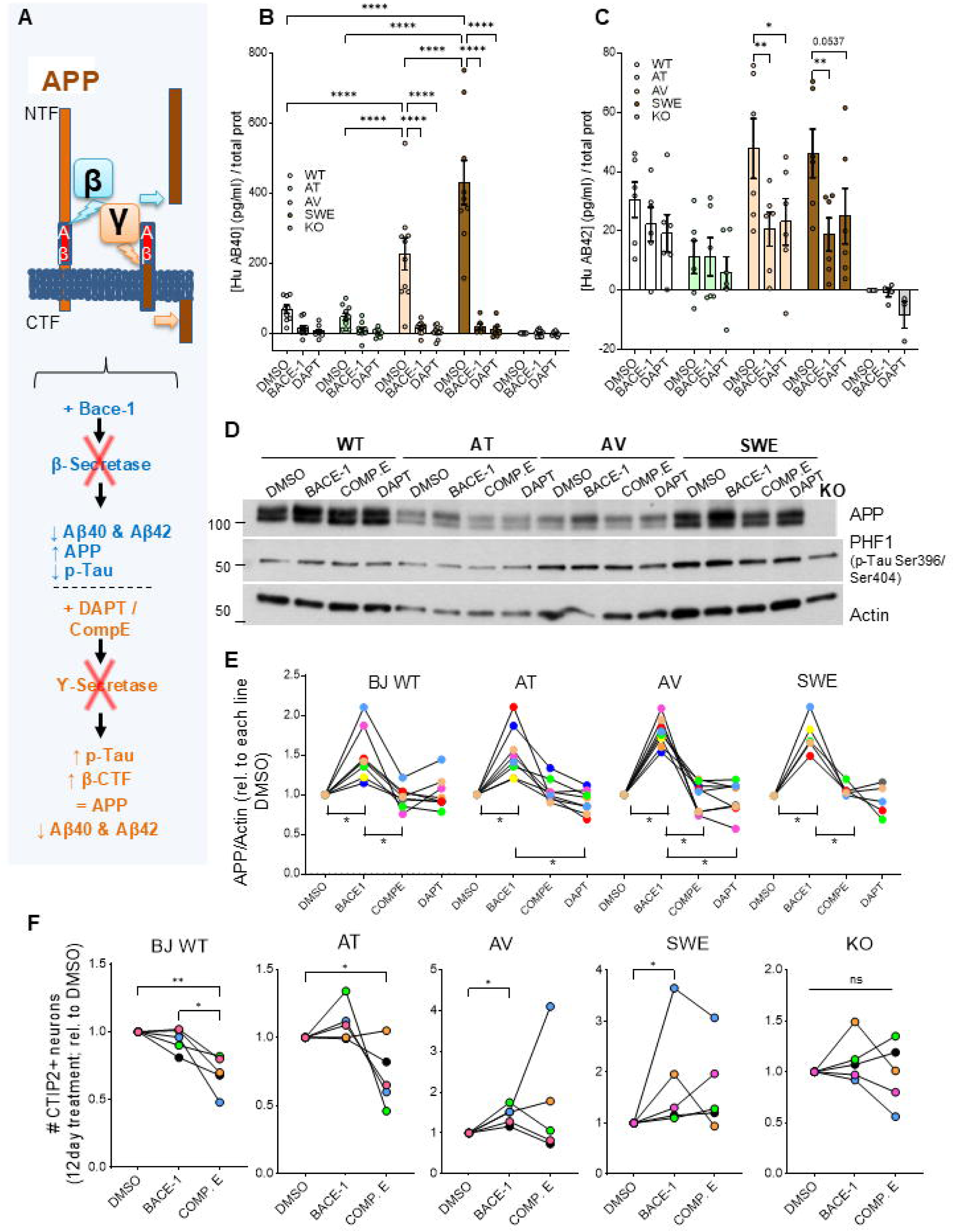
Isogenic familial AD cortical neurons are amenable to treatments that modulate Aß peptide production. (**A**) Schematic representation of *APP* cleavage sites and expected effects of the secretases inhibitors used. ELISA quantification of Aβ40 (**B**) and Aβ42 (**C**) levels in the culture media from fAD CxNs after 12 days in culture treated with the β-secretase inhibitor Bace-1 (5µm) or γ-secretase inhibitors, DAPT and Compound E (5nM). (**D**) Representative western blot from isogenic fAD CxNs treated with the β- or γ-secretase inhibitors for 12 days. *APP*, p-Tau/PHF1 (Ser396/Ser404), total Tau and actin levels are shown and quantification of *APP* levels are shown in (**E**). Values are relative to the DMSO-treated CxNs within each AD line. (**F**) Quantification of the number of CTIP2^+^ neurons for each of the isogenic fAD cultures after treatment with vehicle (DMSO), BACE-1 or Compound E for 12 days. Values shown are relative to the DMSO-treated CxNs within each fAD line (Two-way ANOVA followed by Bonferroni Multiple Comparison Test).

### Familial AD cortical organoids show Aβ aggregation

In recent years, cortical organoids (CxOs) have shown great potential for modeling human neurodevelopment and neurological diseases in vitro ^47^. These organoids have the capacity to self-organize and give rise to highly complex three-dimensional structures and can be maintained in culture for long periods of time, which is often necessary to recapitulate pathological features of late-onset neurodegenerative diseases ^48^. Notably, CxOs possess a degree of structural complexity that enables the formation of an interstitial compartment, a crucial feature for modeling disorders like AD, where pathological protein aggregates accumulate in the extracellular space ^49,50^. We hypothesized that long-term culture of our isogenic AD organoids would facilitate the formation and detection of Aβ aggregates, a hallmark of AD pathology. Following a well-established protocol that involved the generation of stem cell aggregates in ultra-low attachment 96-well plates and subsequent sphere embedding in matrigel ^50^, we generated CxOs and maintained them in culture for up to 180 days (Figure 4A). First, to comprehensively characterize the cellular composition of CxOs and assess whether they recapitulate the diversity of cell types found in the developing human cortex, we performed scRNA-seq on day 45 WT and fAD (AV and SWE) CxOs. This approach is essential for resolving cellular heterogeneity of brain organoid systems and benchmarking in vitro models against in vivo developmental trajectories ^15,51^. Consistent with prior studies, our CxOs contained a rich variety of neural cell types, including apical and outer radial glia, proliferative progenitors, neuroblasts, both immature and mature excitatory and inhibitory neurons (Figure 4B and S3) and choroid plexus cells, These data confirm that our model recapitulates key features of early corticogenesis.

**Figure 4.**
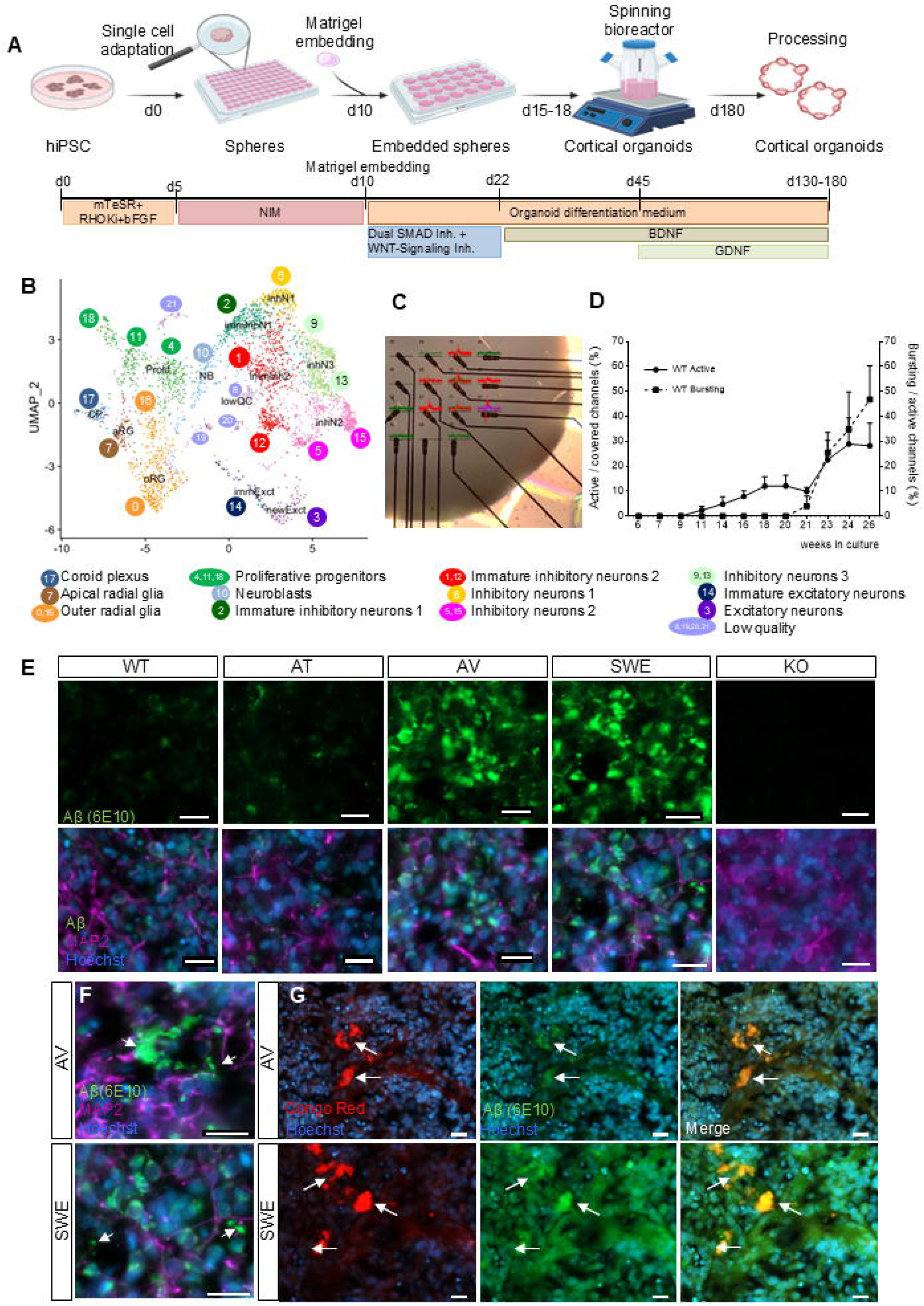
Cortical organoids carrying the fAD-*APP* mutations accumulate extracellular Aβ. **(A)** Schematic representation of the protocol followed to generate cortical organoids (CxOs) from the isogenic fAD hiPSC lines. (**B**) UMAP dimensionality reduction generated from integration of all three samples, only day 45 WT CxOs sample is shown here. Coloring according to identified main clusters. (**C**) Representative image showing a CxO placed on a Multi-electro array (MEA) chip and electrodes recording electrical activity (red). (**D**) Quantification of the percentage of electrodes registering basal (straight line) and bursting activity (dash line, indicator of maturation of neural networks) from WT CxOs being longitudinally recorded using MEAs from week 6 to week 26 of development. (**E**) Representative confocal images from isogenic fAD CxOs fixed after 130 days in culture and immunostained against Aβ with Aβ-6E10 antibody (green), and MAP2 (purple), and nuclei with Hoechst (blue). Scale bar, 20 µm. (**F**) Magnified images from AV and SWE day 130-organoid sections showing Aβ, MAP2 and Hoechst immunostaining. Arrows indicate Aβ-6E10+ deposits. Scale bar, 20 µm. (**G**) Magnified images from AV and SWE 130 day-organoids showing Aβ+ plaques (β-6E10, green) also positive for Congo Red (red) indicated by arrows. Scale bar, 20 µm.

Next, to evaluate the functional maturation and network connectivity of our CxOs, we performed longitudinal multi-electrode array (MEA) recordings on WT organoids from week 6 to week 26 of differentiation. Spontaneous electrical activity emerged around week 11, as shown by a rising percentage of active electrodes relative to those covered by the organoid (Figure 4C-D). Network activity progressively increased through week 26, reflecting ongoing neuronal maturation and synaptogenesis (Figure 4C-D). Notably, bursting activity -defined as trains of high-frequency action potentials, and indicative of functional neuronal networks-emerged consistently from week 20, signaling the establishment of more mature, coordinated network dynamics within the organoids over time (Figure 4C-D). These findings confirm that our CxOs are capable of generating functional, maturing neuronal networks in vitro. Finally, to assess Aβ aggregation, we immunostained CxOs against Aβ and observed prominent intracellular and extracellular Aβ aggregates in both AV and SWE organoids, in stark contrast to WT, AT, and *APP* KO CxOs, where little or no signal was detected (Figure 4E). Staining with the β-amyloid-specific dye Congo Red further confirmed the presence of amyloidogenic deposits (Figure 4F-G). These results demonstrate that our isogenic AD in vitro model recapitulates a fundamental AD hallmark – amyloid aggregation – an outcome that has been difficult to achieve in hiPSC-derived models without overexpression of mutant *APP* and/or *PSEN1* ^52–54^.

### Deleterious *APP* variants preferentially affect distinct molecular pathways

To investigate the potential proteomic changes caused by the introduced SNPs in the *APP* gene underlying the differential CxN survival, we differentiated the isogenic AD lines into CxOs for 45 days and performed multiplexed tandem mass tag (TMT) quantitative mass spectrometry. Nearly 8000 proteins were quantified across three independent differentiation experiments. As expected, principal component analysis revealed minimal variance between lines and individual samples (Figure S4A). Compared to WT, we identified significant changes in protein abundance in all genotypes: 125 proteins in the protective AT line, 87 in AV, 60 in SWE, and 103 in *APP* KO CxOs (Figure 5A and S4B). Interestingly, only five proteins were significantly over- (the lysosomal protease CTSB and the calmodulin regulator protein PCP4) or under-represented (the translation initiation factor 1 (eIF1), a ribonucleoprotein complex component that mediates the translation, SRP19, and alpha-1,3-mannosyltransferase (ALG3), crucial for protein folding and stability) in both fAD lines (Figure 5B). These shared changes suggest partial convergence on pathways affecting translation and lysosomal function, while the largely distinct protein profiles highlight divergent molecular consequences of the two pathogenic mutations. Additionally, 2 proteins showed changes in opposite directions between the fAD CxOs and the ones carrying the protective AT variant: catechol-O-methyltransferase (COMT), central for catecholamine breakdown, being significantly downregulated in SWE while over-represented in AT compared to WT and FAM125A (ESCRT-I complex protein), which was significantly upregulated in AV but downregulated in AT (Figure 5B).

**Figure 5.**
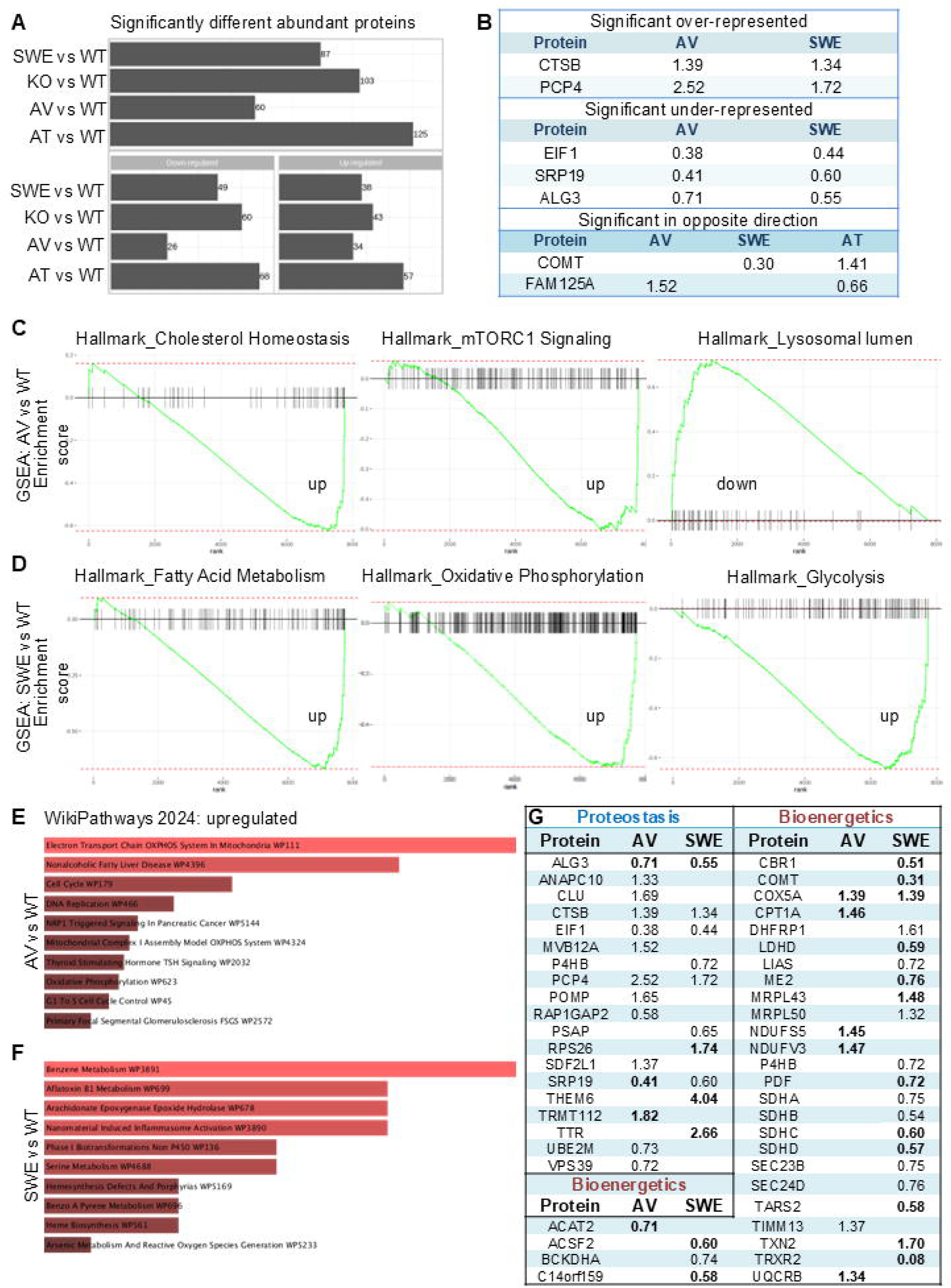
Different deleterious *APP* variants selectively affect distinct molecular pathways. (**A**) Bar chart of global proteomic analysis (TMT-MS) from day45 isogenic fAD cortical spheres showing differential protein abundance between WT, AT, AV, SWE and KO (significance threshold cut-off of p-value <0.05 and fold change ≥1.3). Bottom panels show the number of proteins under-represented (left) and over-represented (right) in the fAD cortical spheres compared to WT. (**B**) Table showing the significantly over- and under-represented proteins shared between AV and SWE cortical spheres compared to WT and over- or under-represented proteins in AV or SWE cortical spheres compared to WT that move in opposite direction in the protected AT cortical spheres (bottom). Gene Set Enrichment Analysis (GSEA) plots performed on proteins ranked based on their relative abundance on day 45 AV (**C**) or SWE (**D**) cortical spheres compared to WT. Three selected pathways significantly enriched for a p-adjust value <0.1 compared to WT are shown. (**E**) Bar plot showing the top 10 enriched pathways in the fAD-AV cortical spheres compared to WT identified by WikiPathways analysis using the protein list specified in A. Pathways ranked by p-value. (**F**) Bar plot showing the top 10 cellular components in the fAD-SWE cortical spheres compared to WT identified by WikiPathways analysis, ranked by p-value. (**G**) Significantly different abundant proteins detected by TMT-MS between fAD-AV or fAD-SWE and WT day45 cortical spheres associated with proteostasis and bioenergetics pathways, respectively.

Gene set enrichment analysis (GSEA) of significantly differentially abundant proteins revealed distinct pathway enrichments. In AV CxOs, cholesterol (Ch) homeostasis, mTORC1 signaling and lysosomal lumen pathways were among the top 5 hallmarks (Figure 5C). In SWE CxOs, fatty acid metabolism, oxidative phosphorylation, and glycolysis were among the top 5 hallmarks for the SWE vs WT set (Figure 5D). All of these (except lysosomal lumen in AV), showed low enrichment scores (ESs), suggesting most of the MS-detected proteins associated with these pathways were upregulated. Conversely, the AT CxOs showed high ESs scores for oxidative phosphorylation and redox homeostasis (Figure S4D) indicating a protective upregulation of mitochondrial resilience pathways. Further pathway analysis confirmed dysregulation of cholesterol metabolism and proteostasis in AV organoids and impaired mitochondrial tricarboxylic acid (TCA) cycle specifically in the SWE (Figure 5E-G and S4C). Interestingly, the most upregulated pathway in the AT CxOs was the metabolism of leucine, isoleucine and valine, collectively known as branched-chain amino acids (BCAAs) and linked to neuroprotective effects ^55,56^ (Figure S4E).

To identify functional relationships among differentially expressed proteins in our isogenic fAD CxOs, we applied STRING network analysis (https://string-db.org/) to the mass spectrometry datasets. This analysis revealed distinct, variant-specific clusters of interacting proteins. In AV organoids, major clusters were associated with proteostasis, the electron transport chain (ETC), the ESCRT I complex, and cholesterol biosynthesis. In contrast, SWE organoids showed enrichment for proteins involved in the TCA cycle and mitochondrial fatty acid β-oxidation, with smaller modules related to polysomal ribosomes and lysosomal function. The protective AT line showed interaction clusters associated with synaptic plasticity and mitochondrial β-oxidation, as well as a minor cluster linked to the actin cytoskeleton. Finally, *APP* KO organoids exhibited enrichment for steroid biosynthesis, TCA cycle components, and proteins involved in synapse and brain development (Figure S5). Altogether, these data support the hypothesis that distinct *APP* variants induce unique, variant-specific proteomic signatures and differentially perturb key cellular pathways relevant to AD pathogenesis.

### Validation of variant-specific dysregulated proteins in familial AD cortical organoids

To validate selected proteomic alterations identified in our isogenic fAD CxOs, we focused on representative markers associated with lysosomal and mitochondrial function, two cellular systems increasingly implicated in early AD pathogenesis. We selected the lysosomal hydrolase Cathepsin B (CTSB) and the ETC subunit SDHB for targeted analysis. CTSB was chosen for its role as a key lysosomal hydrolase implicated in autophagic-lysosomal impairment -a widely acknowledged early feature of AD pathology. Recent studies have shown that compromised lysosomal acidification, increased pro-CTSB and CTSB-positive lysosomes, and upregulation of lysosomal gene transcripts – indicative of enhanced lysosomal biogenesis – are among the earliest detectable changes in both human AD brains and mouse models. These precede plaque formation and gliosis and may contribute to downstream neurodegeneration ^57^. Western blotting of day 45 CxO lysates confirmed a significant increase in CTSB levels in AV CxOs compared to WT, AT and SWE CxOs, in both the larger precursor form, pro-CTSB, and the mature protein (Figure 6A-B), corroborating our MS findings. Interestingly, an upward trend for CTSB abundance was also observed in *APP* KO CxOs.

**Figure 6.**
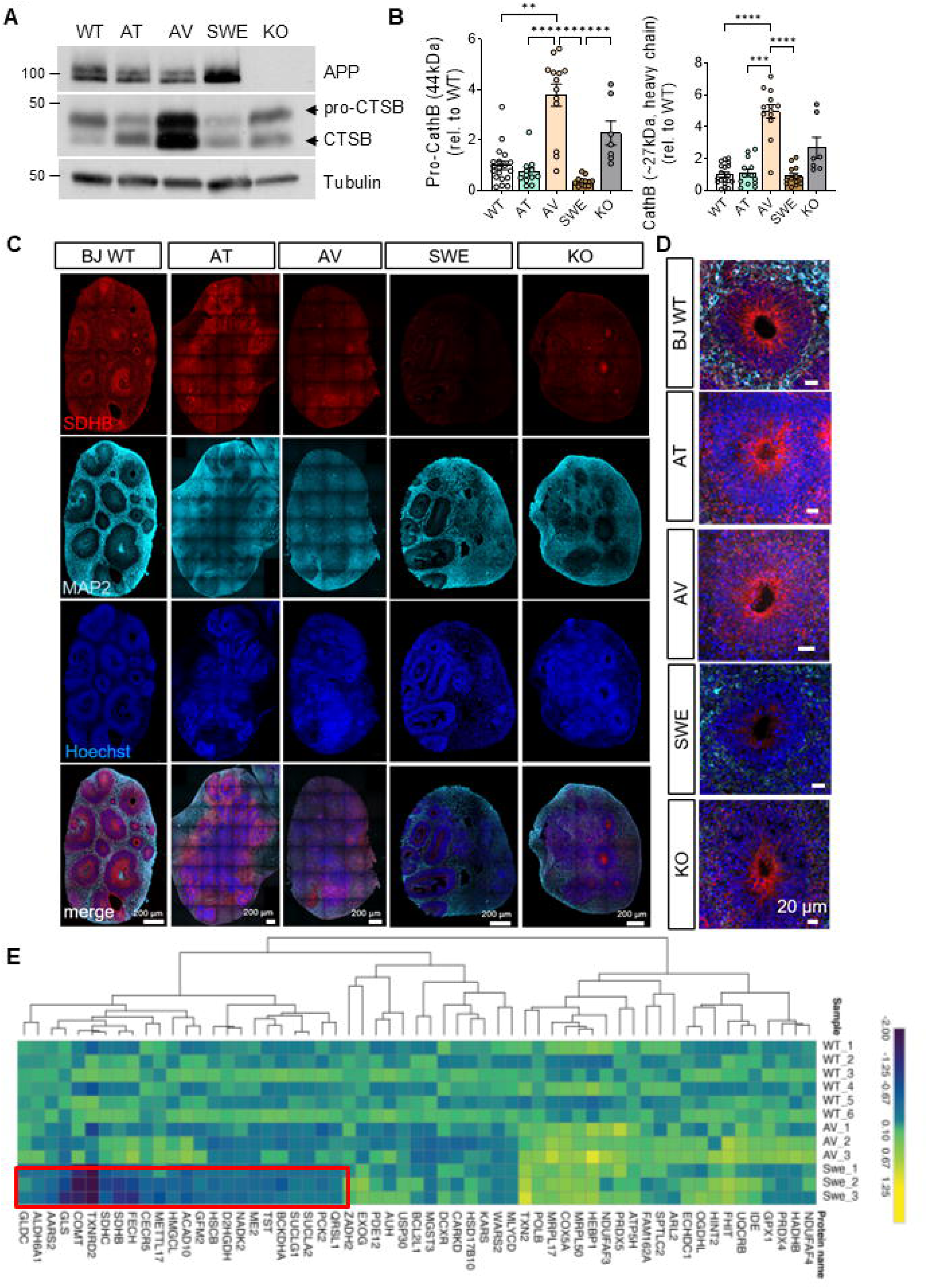
A673V mutation affects lysosomal biology, whereas KM670/671NL influences mitochondrial bioenergetics. (**A**) Representative immunoblot of the lysosomal hydrolase Cathepsin B (CTSB) and the chaperone Clusterin (CLU) found significantly over-represented in the fAD cortical spheres by TMT-MS and quantification (**B**). (**C**) Representative confocal images from day 45 CxOs cryosectioned and immunostained against the mitochondrial protein SDHB (part of complex II of the electron transport chain) (red) and MAP2 (cyan), nuclei are labeled with Hoechst (blue). Scale bar, 200 µm. Magnification of a ventricle-like structure from the immunostained CxOs are shown in (**D**). (**E**) Heatmap analysis showing mitochondrial function or localization associated proteins with differential abundance in AV or SWE day 45 CxNs compared WT detected by TMT-MS and aligned to MitoCarta 3.0. (Cut-offs applied: minimum 2 peptides/protein quantified, intra-group CV<30%, ANOVA and p-value<0.05).

SDHB, a key component of complex II in ETC, was selected due to its central role in linking TCA cycle activity with oxidative phosphorylation. Immunostaining against SDHB revealed markedly reduced signal in day 45 SWE CxO, both throughout entire sections and specifically in areas corresponding to ventricle-like structures negative for the pan-neuronal marker MAP2 (Figure 6C-D). This is consistent with the predominance of mitochondrial proteins among dysregulated MS hits in the SWE MS dataset (Figure 5G). Indeed, over 60% of significantly altered proteins in SWE CxOs map to MitoCarta 3.0, indicating a broad impact on mitochondrial metabolism (Figure 6E). This focus aligns with prior studies demonstrating mitochondrial dysfunction in SWE *APP* models, including APP’s interaction with mitochondrial chaperone HSP60 and its mislocalization to mitochondria, contributing to oxidative stress and impaired respiration in both AD patient tissue and transgenic mice ^58,59^. To validate our isogenic fAD CxOs findings, we analyzed the expression of two key mitochondrial proteins – SDHB and TRXR2 (TXNRD2) – identified in SWE CxOs, in dorsolateral prefrontal cortex (DLPFC) sections from two AD patients and two age-matched controls. Confocal imaging revealed significantly decreased SDHB and TRXR2 puncta intensity in AD samples versus controls (Figure 7A-C), while nuclear density (Hoechst-labeled cells) remained unchanged (Figure 7D), supporting genuine mitochondrial impairment independent of neuronal loss or structural atrophy. Altogether, these data corroborate our proteomic findings and are consistent with a model in which distinct *APP* mutations drive divergent early pathogenic mechanisms in fAD: the AV mutation skews toward early lysosomal/proteostasis dysfunction, while SWE mutation predominantly impairs mitochondrial bioenergetics. Our findings thus underscore the heterogeneity of early pathogenic cascades in fAD and highlight specific vulnerabilities linked to individual *APP* variants.

**Figure 7.**
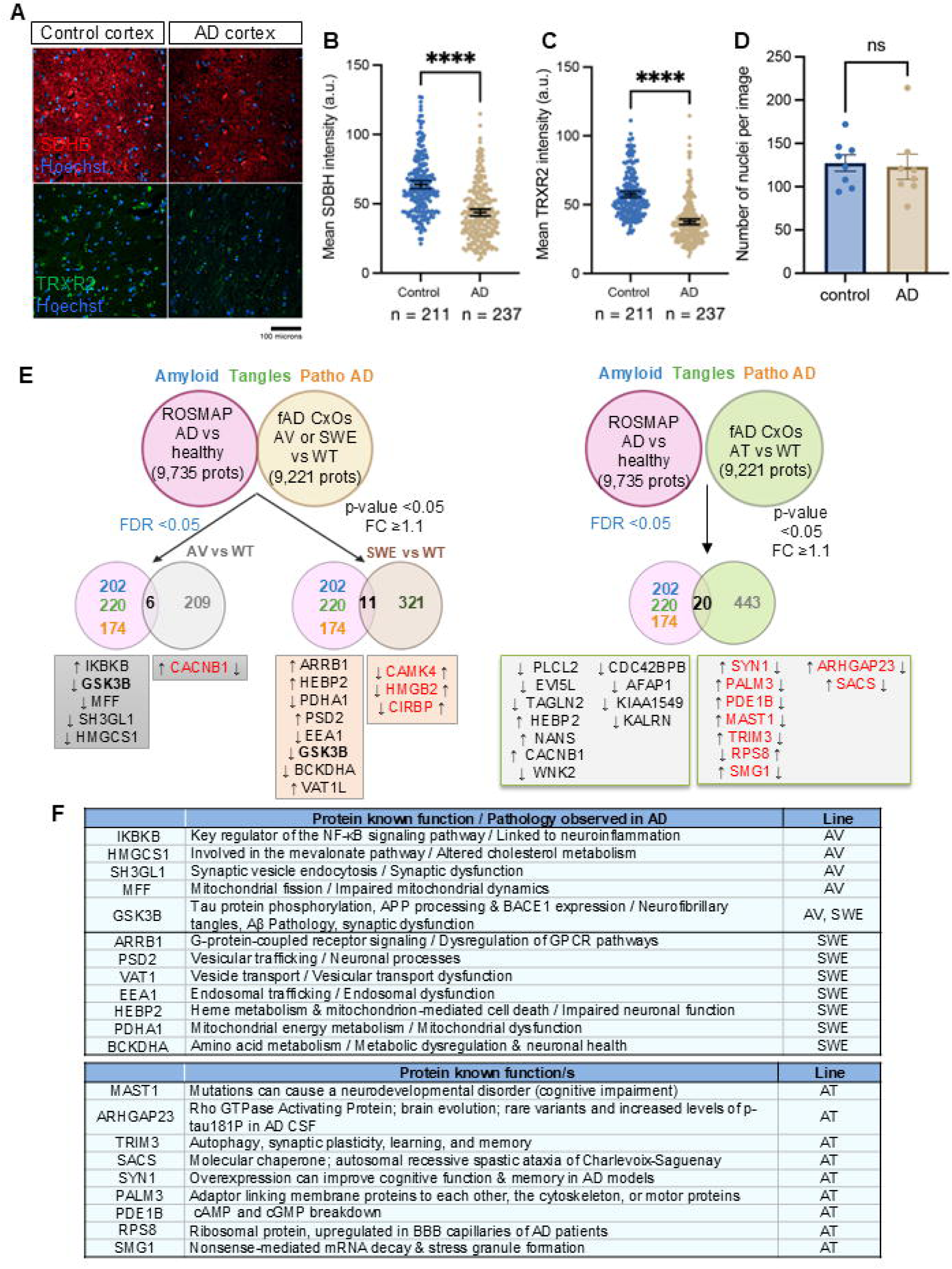
Comparison of MS analysis from fAD-*APP* CxOs with post-mortem AD brains (ROSMAP) reveals new players potentially contributing to AD pathology. (**A**) Representative confocal images from post-mortem AD brain sections (dorsolateral prefrontal cortex (DLPFC)) immunostained against SDHB (Red) and TRXR2 (green). Nuclei are labeled with Hoechst. Quantification of the mean intensity of SDHB+ puncta (**B**) and TRXR2+ puncta (**C**) from two healthy control and two AD DLPFC brain regions (“n” indicates the number of individual cells automatically identified by Hoechst proximity and quantified by the image analysis software). (**D**) Quantification of the number of Hoechst+ nuclei per quantified image. (**E**) Schematic representation of the computational approach followed to compare the human ROSMAP MS data with the MS from our fAD CxOs (left, ROSMAP vs AV and SWE; right, ROSMAP vs AT CxOs). Proteins found significantly under- or over-represented in both datasets are indicated in black. Proteins found dysregulated in opposite directions between both datasets are indicated in red (left arrows refer to the change observed in the post-mortem AD brains vs healthy controls and right arrows refer to the change observed in the CxOs vs BJ WT). (**F**) Tables showing main known functions for the proteins found moving in opposite directions between both MS datasets for AV/SWE (top) and AT (bottom).

### Cross-model comparison identifies conserved proteomic alterations in fAD organoids and human AD brains

To determine whether the proteomic changes observed in our isogenic fAD organoids are conserved in human AD, we compared the AV and SWE CxO proteomes with post-mortem dorsolateral prefrontal cortex (DLPFC) proteomic data from the ROSMAP cohort, available through the AMP-AD portal. This cross-model alignment revealed six and eleven proteins with significantly altered abundance in both AV and SWE CxOs and AD brain tissue, respectively, with most changes following the same directionality across systems (Figure 7E). Notably, several of these overlapping proteins relate to cholesterol metabolism, vesicle trafficking, mitochondrial dysfunction, and neuroinflammation, pathways that we previously linked to distinct *APP* mutations. However, their contributions to AD pathology have remained largely underexplored, likely due to bulk tissue averaging, late disease stages, or limited proteomic depth in post-mortem samples. Among concordantly downregulated proteins in AV CxOs and AD brains was HMGCS1, a key enzyme in the mevalonate pathway that regulates cholesterol biosynthesis. Disruption of cholesterol homeostasis has been linked to synaptic dysfunction ^60^. SH3GL1 (Endophilin A2), essential for clathrin-mediated synaptic vesicle endocytosis, was also reduced in AV CxOs and AD brains, consistent with impaired vesicle cycling reported in AD models ^61^. In SWE CxOs and AD brains, we observed reduced levels of PDHA1, the E1-alpha subunit of pyruvate dehydrogenase, and BCKDHA, involved in branched-chain amino acid catabolism, implicating dysregulation at the glycolysis–TCA interface and branched-chain amino acid (BCAA) metabolism, respectively, both recently linked to microglial and metabolic dysfunction in AD ^62–64^. Interestingly, GSK3B, a kinase central to tau phosphorylation and synaptic toxicity in AD, was elevated across both AV and SWE CxOs and AD brains, further validating its pathophysiological relevance (Figure 7E-F).

In contrast, comparison between the protective AT CxOs and ROSMAP AD brains highlighted twenty shared dysregulated proteins, approximately half of which showed inverse regulation, being upregulated in AT organoids but downregulated in AD brains or vice versa (Figure 7E-F). We postulate that proteins showing opposing dynamics in AT vs WT and ROSMAP AD may represent protective factors. Among these was MAST1 (Microtubule-associated serine/threonine kinase 1), which is involved in neuronal morphogenesis and synaptic stability. Although underexplored in AD, kinase signaling through MAST1 has been implicated in cognitive deficits and could influence neuronal resilience^65^. ARHGAP23, a Rho GTPase-activating protein, also showed opposite regulation. Interestingly, several proteins in the ARHGAP family (e.g., ARHGAP35 and ARHGAP5) have been linked to brain development and evolution ^66^, and others (e.g. ARHGAP22 and ARHGAP10) have also been associated with synaptic function, learning, and memory ^67^, suggesting a potential neuroprotective function. Other candidates such as TRIM3, part of the tripartite motif family involved in ubiquitination, autophagy and synaptic plasticity ^68^, were also downregulated in AD brains but upregulated in AT organoids, highlighting its potential role in conferring resilience in the AT context.

Overall, these data indicate that our fAD CxOs recapitulate core aspects of AD-related proteomic dysregulation seen in human brains, particularly involving proteostasis and mitochondrial deficits. Importantly, the detection of oppositely regulated proteins in the protective AT line unveils candidate factors that may mediate resilience and disease modification. This cross-model comparison highlights the translational utility of isogenic organoids for uncovering early, mutation-specific AD drivers and therapeutic targets, while disentangling them from later-stage, non-causal changes observed in human brain tissue.

### Precise modulation of variant-specific pathways selectively ameliorates neuronal death

To investigate whether the mutation-specific vulnerabilities observed in our isogenic fAD CxOs are amenable to therapeutic intervention, we tested two pathway-specific approaches based on our proteomic findings and phenotypic validations. First, since AV-mutant CxOs exhibited pronounced dysregulation of proteostasis and lysosomal function, we overexpressed TFEB, a master regulator of lysosomal biogenesis and autophagy ^69,70^ (Figure 8A). TFEB has previously been shown to ameliorate AD-like pathology in multiple in vivo models, including *APP*/PS1 ^71,72^ and 5XFAD mice ^73^, by enhancing lysosomal clearance and reducing Aβ accumulation. More recently, it has also been shown to promote lysosomal homeostasis and microglia activation in a mouse model of tauopathy ^74^. Consistent with our earlier findings (Figure 1C-F), we observed that CxNs from AV, SWE, and *APP* KO lines exhibited increased cell death relative to WT, whereas neurons carrying the protective AT variant were more resilient (Figure 8B-C). Upon lentiviral overexpression of TFEB, AV CxNs showed a significant increase in CTIP2+ neuron survival compared to scramble controls, while no such effect was seen in SWE, AT, WT, or KO neurons (Figure 8B-C). This suggests that TFEB activation specifically rescues proteostasis-related vulnerability in AV neurons. Next, we targeted the mitochondrial dysfunction observed in SWE CxOs, where proteomic analyses revealed altered oxidative phosphorylation and TCA cycle activity. As impaired ETC function can elevate reactive oxygen species (ROS) levels, we hypothesized that SWE neuron death might be mediated by ferroptosis, an iron- and ROS-dependent form of regulated cell death increasingly implicated in AD pathogenesis ^75,76^ although the mechanism remains unclear. Treatment with either Liproxstatin-1 or Ferrostatin-1 -well-established inhibitors of lipid peroxidation and ferroptosis^77^- led to a clear rescue of CTIP2^+^ neuronal survival in SWE cultures, with no positive effect in WT, AT, AV or KO cultures lines (Figure 8D-F).

**Figure 8.**
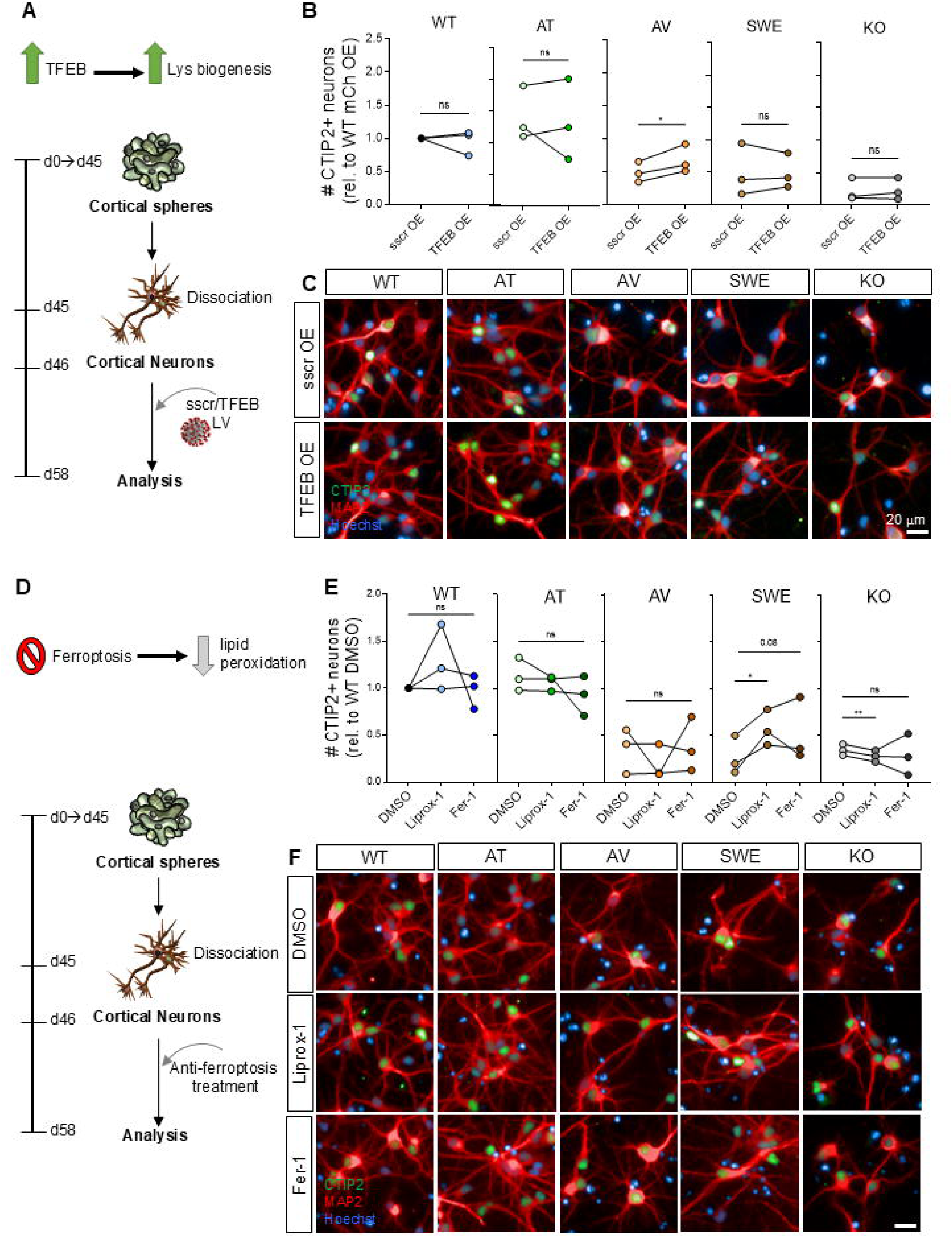
Targeting of *APP* mutation-specific molecular mechanisms rescues neuronal death in a selective manner. (**A**) Schematic representation of the experimental design followed to boost TFEB levels in CxN cultures. (**B**) Quantification of the number of CTIP2+ CxNs dissociated from day45 fAD isogenic cortical spheres and transduced with either a lentivirus (LV) carrying a scramble control (sscr OE) or TFEB CDS (TFEB OE). Values are shown relative to the BJ WT CxNs transduced with the sscr LV. (Two-tailed Paired t-test). (**C**) Representative confocal images showing CxN cultures treated with sscr or TFEB LVs and fixed and immunostained 12 days later against CTIP2 (green) and MAP2 (red). Nuclei are labeled with Hoechst. Scale bar, 20 µm. (**D**) Schematic representation of the experimental design followed to reduce lipid peroxidation by inhibiting the ferroptosis pathway in CxN cultures. (**E**) Quantification of the number of CTIP2+ CxNs dissociated from day45 fAD isogenic cortical spheres and treated with the ferroptosis inhibitors Liproxstatin-1 (Liprox-1, 1 μM) or Ferrostatin-1 (Fer-1, 1 μM), or DMSO as vehicle control. Values are shown relative to the BJ WT CxNs treated with DMSO. (One-way ANOVA followed by Fisher’s LSD test). (**F**) Representative confocal images showing CxN cultures treated with DMSO, Liprox-1 or Fer-1 for 12 days, fixed and immunostained against CTIP2 (green) and MAP2 (red). Nuclei are labeled with Hoechst. Scale bar, 20 µm.

Together, these results demonstrate that variant-specific pathways identified in fAD organoids are functionally actionable, and that targeted interventions, such as TFEB activation or ferroptosis inhibition, can selectively restore neuronal survival in a mutation-dependent manner.

## Discussion

By generating a panel of isogenic hiPSC lines carrying pathogenic or protective *APP* variants and differentiating them into cortical organoids and their derived neurons, we showed that early neuronal vulnerabilities can be selectively reversed through pathway-specific interventions. This study supports the feasibility of presymptomatic therapeutic targeting in AD and reinforces the promise of a precision-medicine framework in which treatments are matched to the distinct pathogenic trajectories initiated by individual *APP* variants. We also demonstrate the power of isogenic cortical organoid systems and the analytical tools they enable to uncover variant-tailored therapeutic strategies for genetically distinct forms of fAD.

AD emerges through a long, clinically silent period during which selective neuronal populations experience progressive molecular stress well before overt neurodegeneration. Identifying the earliest human-relevant events in this preclinical window has been limited by reliance on postmortem tissue and animal models that incompletely reproduce human APP processing, cortical maturation, and transcriptional programs ^13,78^. Here, we used an isogenic panel of hiPSC-derived cortical neurons and organoids carrying pathogenic (A673V, Swedish) and protective (A673T) *APP* mutations to define early, variant-encoded molecular vulnerabilities in a controlled human context. Our findings reveal that distinct fAD mutations do not collapse immediately onto a single pathway but instead generate discrete, targetable early states centered on proteostasis/lysosomal dysfunction (A673V) or mitochondrial bioenergetics failure (Swedish). Notably, extended culture enabled the formation of extracellular Aβ deposits without *APP/PSEN1* overexpression, demonstrating that physiologic expression of patient-relevant *APP* mutations is sufficient to induce hallmark AD pathology in 3D. These findings underscore the utility of developmentally appropriate human organoids for modeling early-stage AD and for designing interventions in a biologically relevant context.

Despite comparable increases in β-secretase processing across pathogenic lines, the downstream molecular consequences diverged sharply. In A673V organoids, proteomic analyses uncovered broad upregulation of lysosomal hydrolases, ESCRT-I components, and cholesterol-associated factors, consistent with early lysosomal and proteostasis stress. These findings align with recent work showing that A673V-modified C99 is preferentially degraded by the proteasome while mature A673V APP is more efficiently targeted to lysosomes ^29^, suggesting that this mutation imposes a dual burden on degradative systems. In contrast, Swedish organoids displayed widespread perturbations in the tricarboxylic acid (TCA) cycle, oxidative phosphorylation, and mitochondrial redox pathways. This mitochondrial vulnerability resonates with prior studies reporting that Swedish APP can perturb APP–mitochondria interactions, elevate ROS, disrupt respiratory chain activity, and impair mitophagy in mouse and human systems^58,79–83^, and our data place these alterations as a prominent early feature in an isogenic human neural context.

These divergent downstream consequences may also reflect fundamental biochemical differences between the Aβ species produced by each mutation. Whereas the Swedish variant increases total production of wild-type Aβ40/42, the A673V mutation generates an N-terminally modified Aβ peptide (A2V), which alters oligomerization kinetics and fibrillogenic potential ^22,24^. It is therefore plausible that qualitative differences in Aβ chemistry, not only quantitative differences in Aβ burden, bias neurons toward distinct stress responses. In this framework, A673V-modified Aβ/Aβ-C99 species may preferentially overwhelm proteasomal and lysosomal pathways, whereas elevated levels of wild-type Aβ in Swedish neurons may exert stronger pressure on mitochondrial metabolism and redox balance. Although our current dataset cannot resolve causality, and confirmation would require additional isogenic lines carrying *APP* mutations outside the Aβ sequence, this model provides a testable hypothesis linking mutation-specific Aβ biochemistry to the early, divergent vulnerabilities we observe. This framework may also help explain clinical and molecular heterogeneity observed in AD patients, in whom early lysosomal or mitochondrial dysfunction often appears unevenly distributed across individuals and brain regions.

These mutation-specific profiles intersect with an enduring question in the AD field: do lysosomal/autophagic defects or mitochondrial dysfunction arise first? While some studies position lysosomal failure as the earliest detectable abnormality in human AD brain ^46,57^, others highlight early mitochondrial ROS, metabolic reprogramming, and Aβ-induced alterations in mitochondrial dynamics ^84,85^. Rather than supporting a strict hierarchy, our isogenic system suggests an integrative model in which *APP* mutations bias neurons toward distinct entry points along a shared lysosome–mitochondria stress circuit. Indeed, mounting evidence implies mitochondrial-lysosomal crosstalk in AD, where proteostasis failure, such as lysosomal membrane permeabilization, compromises mitochondrial integrity, and conversely, mitochondrial oxidative stress can collapse lysosomal acidification and autophagic flux ^57,86–89^, while mitochondrial oxidative stress can. Within this framework, either arm of the circuit may serve as the initial point of failure depending on genetic context. Although our cross-sectional analyses cannot establish temporal precedence, this model accommodates both “proteostasis-first” and “mitochondria-first” views while providing a mechanistic explanation for the divergent states induced by A673V and Swedish mutations. Crucially, these early states are selectively reversible. Enhancing lysosomal function via TFEB overexpression rescued neuronal survival only in A673V cultures, whereas Swedish neurons responded uniquely to inhibition of lipid peroxidation and ferroptosis. These highly specific rescue patterns argue against the idea that the two mutations simply represent earlier and later positions along a single linear trajectory; if that were the case, some overlap in responsiveness would be expected. Instead, each mutation produces a distinct, mutation-encoded vulnerability that defines the earliest actionable molecular state in human cortical neurons. The specificity of these rescue responses suggests that identifying early, variant-defined cellular states may be key to designing interventions that are effective before irreversible neurodegeneration occurs.

Comparisons with the ROSMAP postmortem proteome reinforce the translational relevance of these pathways. Both mutations overlapped with AD-associated dysregulation in cholesterol biosynthesis (HMGCS1), vesicle trafficking (SH3GL1), mitochondrial metabolism (DLD, BCKDHA), redox regulation (TRXR2), and the tau kinase GSK3B. In contrast, the protective A673T variant demonstrated inverse regulation of several AD-dysregulated proteins, including TRIM3, ARHGAP23, and MAST1, highlighting candidate resilience mechanisms that become visible only in a genetically matched human model. It is worth noting that several mutation-specific mitochondrial and redox-related proteins highlighted in our organoid analyses, including SDHB and TRXR2, were not detected in the ROSMAP bulk proteomic dataset. This discrepancy likely reflects technical rather than biological differences, as mitochondrial and other low-abundance proteins are often underrepresented in mass-spectrometry datasets derived from heterogeneous postmortem tissue, where cell-type–specific signals are diluted by bulk cortical homogenates. In contrast, organoid proteomic profiling captures pathway alterations emerging within a defined neuronal lineage, enabling the detection of early, mutation-specific vulnerabilities that may be masked in large-scale human datasets. Consistent with this interpretation, immunostaining of human AD brain samples confirmed altered abundance of both SDHB and TXNRD2, reinforcing the physiological relevance of the organoid-derived signatures.

Overall, our findings position isogenic cortical organoids as a powerful platform for mechanistically dissecting the earliest, mutation-specific perturbations in human neurons. By revealing that A673V and Swedish mutations generate divergent, targetable molecular states, rather than a single unified trajectory, this work reframes fAD mutations as precision tools for understanding mechanistic heterogeneity at AD onset. More broadly, these results illustrate the strength of stem-cell–derived human neural models for resolving early pathogenic programs that cannot be accessed in postmortem tissue or recapitulated in animals. Mapping how specific genetic variants bias neurons toward defined lysosomal or mitochondrial stress states provides a foundation for precision therapeutic strategies targeting the earliest, and potentially reversible, phases of AD pathogenesis. As such, this work lays a foundation for systematically mapping how distinct genetic variants shape early and longitudinal actionable phases of AD pathogenesis and for leveraging human organoid systems to accelerate precision therapeutic discovery.

We also note some limitations of our study. We focused on one *APP* variant that does not alter the Aβ sequence (Swedish), but other non–Aβ-sequence mutations near the β-secretase site remain unexplored. Whether these additional variants also produce mitochondrial defects or engage similar early pathogenic pathways will require future investigation. Additionally, although cross-model comparisons with ROSMAP brain proteomic datasets highlight clear concordances, differences in tissue composition, disease stage, and analytical depth limit direct quantitative comparisons. Finally, because our study focuses on familial *APP* variants, the extent to which these variant-specific pathways generalize to late-onset sporadic AD remains uncertain. However, several of the dysregulated pathways we identify, particularly those involving lysosomal, mitochondrial, and cholesterol metabolism, overlap with alterations observed in sporadic AD cohorts. This suggests that while not all mechanisms will extrapolate directly, the variant-defined cellular states may point to broader principles of early AD pathogenesis. Future work integrating single-cell multiomics, longer-term maturation, and multicellular organoid systems incorporating glia and vasculature will be informative to refine the mechanistic and translational insights enabled by our platform.

## Figures and figure legends

**Supplemental Figure 1. Generation of isogenic familial AD hiPSC lines targeting the *APP* gene.** (**A**) Knock-in mutagenesis workflow to generate hiPSC lines carrying *APP* variants. (**B**) Scheme of HR120-PA-AD targeting vector used for knock-in mutagenesis to generate HR120-PA-AT, HR120-PA-AV and HR120-PA-SWE hiPSC lines. (**C**) Isogenic hiPSC lines generated and used for the study. (**D**) Sanger-sequencing results from the successfully targeted fAD clones aligned to the unedited (WT) line showing the regions of interest of the *APP* locus. Two confirmed clones for each line are shown, from top to bottom: AT-line (GCA->ACA), AV-line (GCA->GTA), SWE-line (AAG,ATG->AAT,CTA) and KO line (32 bp deletion). The introduced silent variants in the homology arms to avoid rebinding of the sgRNA and potential subsequent cutting from Cas9 are highlighted in light blue. The sgRNA used in each case is shown as a green arrow in between the reference WT and the targeted clone sequences. (**E**) Representative confocal images from control (WT), AT (carrying protective variant), AV, SWE (carrying pathogenic mutations) and *APP* KO (KO) hiPSCs fixed and immunostained against the pluripotency markers OCT4 (red) and SOX2 (green). Nuclei are stained with Hoechst (blue). Scale bar, 10 µm. Quantification of the percentage of hiPSC nuclei positive for OCT4 (**F**) and SOX2 in (**G**). One-way ANOVA followed by Tukey’s Multiple Comparison Test. (**H**) Western blot from isogenic fAD hiPSC lysates showing *APP* protein levels and quantification (**I**).

**Supplemental Figure 2. Generation and validation of CTIP2:mNeonGreen reporter isogenic fAD lines (associated to Figure 1).** (**A**) Genome targeting scheme designed to introduce the mNeonGreen fluorescent reporter downstream of the CTIP2 locus in the fAD isogenic hiPSC lines. (**B**) Schematic view of the targeting vector used to generate the CTIP2 reporter lines, HR120-PA-CTIP2-P2A-mNeoNGreen. mNeonGreen expression from genome-edited WT cortical spheres after 35 days in culture (**C**) and from CxNs right after day 45 sphere dissociation and plating (**D**). (**E**) Representative images from the CTIP2:mNeonGreen targeted WT hiPSC line showing CxNs immunostaining against CTIP2 (red) and the green signal from the CTIP2 genetically modified locus. Nuclei are stained with Hoechst (blue). Scale bar, 20 µm. (**F**) Quantification of the percentage of CTIP2+ (antibody-based); CTIP2:mNeonGreen double-positive CxNs. (**G**) FACS plots showing the percentage of CTIP2+ neurons from freshly dissociated cortical spheres at the endpoint of the differentiation protocol (∼ day 45). The GFP+ cells indicated in the plots correspond to the CTIP2:mNeonGreen+ CxNs from the reporter isogenic fAD hiPSC lines. The percentage of CTIP2:mNeonGreen+ CxNs for all lines is quantified in (**H**).

**Supplemental Figure 3 (associated to Figure 4).** (**A**) UMAP dimensionality reduction generated from integration of all three samples, day 45 AV and SWE CxOs samples are shown (See Figure-4B for WT sample). Coloring according to identified main clusters. (**B**) Dotplot showing expression levels of key marker genes for the identified cell clusters of day 45 CxOs. (**C**) Feature Plots representation of day 45 WT CxOs highlighting the expression of marker genes for the main identified cell clusters.

**Supplemental Figure 4. TMT mass spec from day 45 cortical spheres derived from the isogenic fAD hiPSC lines (associated to Figure 5).** (**A**) Principal component analysis (PCA) of day 45 isogenic AD cortical spheres subjected to mass spec. (**B**) Heatmap after applying default clustering and row scaling across AV and SWE samples (left) or AT (right) compared to WT using proteins that were quantified with at least two unique peptides, showed a CV<30% and p-value<0.01. Columns represent samples, rows represent proteins, and color intensity represents column Z score, where yellow indicates enriched and blue depleted proteins. (**C**) Bar plot showing the top 10 enriched downregulated pathways in the fAD-AV (top) and fAD-SWE (bottom) cortical spheres compared to WT identified by WikiPathways analysis using the protein list specified in Fig. 5A. Pathways ranked by p-value. (**D**) Gene set enrichment analysis (GSEA) plots performed on proteins ranked based on their relative abundance from day 45 AD-AT cortical spheres compared to WT. Three selected pathways (of the top 6) showing a significantly different enrichment score to WT are shown. **(E**) Bar plot showing the top 10 enriched up- (top) and downregulated (bottom) pathways in the AT cortical spheres compared to WT identified by WikiPathways analysis using the protein list specified in Fig. 5A. Pathways ranked by p-value.

**Supplemental Figure 5.** MS String Clustering Analysis for AV, SWE, AT and *APP* KO lines (Protein–protein interaction networks were analyzed using STRING v12.0 (Szklarczyk et al., 2023), selecting Homo sapiens as the species. Proteins identified by mass spectrometry as significantly regulated (fold change ≥ 1.3 or ≤ −1.3; p < 0.05) were submitted for clustering. The analysis was based on the full STRING network, incorporating both functional and physical protein associations, with a minimum interaction score of 0.400 (medium confidence). Clustering was performed using the Markov Cluster Algorithm (MCL) with an inflation parameter of 1.6. Clusters containing four or more proteins were visually highlighted and annotated. Node colors indicate fold change (red = upregulated, blue = downregulated), and dotted lines represent edges between distinct clusters).

**Supplemental Table 1.** ssOligos ordered to generate the different CRISPR for generating the isogenic fAD hiPSC lines.

**Supplemental Table 2.** Quantified proteins along with their associated TMT reporter ion ratios used for quantitative analysis.

## Materials and methods

### Study approval

The use of the hiPSC lines in the Rodriguez-Muela lab at DZNE was approved by the Ethics Commission at the Technische Universität, Dresden (SR-EK 80022020). The information regarding both the healthy control lines is enclosed in ^41^ and ^90^.

### Maintenance of hiPSCs

hiPSCs were cultured on Matrigel-coated plates (ESC-qualified; BD Biosciences) in mTeSR Plus medium (STEMCELL Technologies) supplemented with 1% (vol/vol) penicillin–streptomycin (Life Technologies). Cells were passaged using ReLeSR (STEMCELL Technologies, 100–0484) and maintained at 37 °C in a humidified incubator with 5% CO₂ and 95% relative humidity, as previously described ^91^.

### Generation of isogenic fAD lines

To generate the APP knockout (KO) line, we employed a single-vector CRISPR–Cas9 approach. The px458 plasmid (Addgene #48138), which expresses both Cas9 and a single-guide RNA (sgRNA), was used to introduce double-strand breaks at exon 2 of the *APP* locus (Supplemental Table 1). Complementary single-stranded DNA oligonucleotides (IDT), including BbsI-compatible overhangs, were annealed and cloned into px458 via BbsI digestion and T7 DNA ligase–mediated ligation in a one-step digestion–ligation reaction. Correct insertion of the sgRNA into px458, yielding the construct pTG-Cr-KO, was confirmed by Sanger sequencing (Microsynth). To generate isogenic knock-in lines carrying familial Alzheimer’s disease–associated and protective APP variants, we employed a two-vector CRISPR–Cas9 targeting strategy adapted from a previously described method. Using the same cloning strategy as above, complementary oligonucleotides encoding sgRNAs targeting exon 16 of the *APP* locus (Supplemental Table 1) were cloned into px458, generating the constructs pTG-Cr-AT, pTG-Cr-AV, and pTG-Cr-SWE. Correct insertion of each sgRNA sequence was verified by Sanger sequencing (Microsynth). For homology-directed repair (HDR), a second plasmid - a modified version of the HR120-PA1 vector (System Biosciences) - served as the donor template. This vector carried the desired single-nucleotide variants (SNVs) and was used to direct HDR following Cas9-mediated cleavage. To reduce vector size, the copGFP, WPRE, and SV40 promoter elements were removed by digestion with EcoRI and NruI. The final donor vector additionally contained a dual-selection cassette (mRuby–T2A–puromycin) flanked by loxP sites to enable preselection of correctly targeted cells. For each knock-in construct, two synthetic double-stranded DNA fragments (gBlocks; IDT) were designed, each comprising 420 bp of homology to the 5′ or 3′ flanking region of APP exon 16. These included the desired SNV and additional silent mutations to prevent Cas9 re-cutting. The 5′ homology arm was inserted into the modified HR120-PA1 backbone via NheI digestion followed by Gibson Assembly (NEB). The 3′ homology arm was subsequently introduced by BamHI digestion and a second round of Gibson Assembly. Integration of both homology arms was verified by Sanger sequencing. hiPSCs were nucleofected using the 4D-Nucleofector System (AMAXA) with the P3 Primary Cell Kit (Lonza) according to the manufacturer’s instructions. Successfully targeted cells were selected with 1 μg/mL puromycin (Life Technologies, A1113802) for 7 days. To excise the selection cassette, cells were nucleofected with pCAG-Cre:GFP (Addgene #13776), and 24 h later, GFP-positive and mRuby-negative cells were enriched by fluorescence-activated cell sorting (FACS). Post-Cre–excised hiPSCs were plated at clonal density and screened for successful HDR and correct insertion using primers flanking the targeted exon. At least two independent clones per line were expanded for downstream experiments.

### Generation of CTIP2:mNeonGreen reporter lines

To generate CTIP2:mNeonGreen isogenic knock-in reporter lines, we employed a two-vector CRISPR–Cas9 targeting strategy. Using the same cloning approach as described above, complementary oligonucleotides encoding sgRNA sequences (Supplemental Table 1), designed to direct Cas9-mediated cleavage near the *CTIP2* stop codon, were cloned into px458, yielding the construct pTG-Cr-CTIP2:mNeonGreen. Correct sgRNA insertion was confirmed by Sanger sequencing (Microsynth). The donor plasmid was derived from a modified version of the HR120-PA1 backbone (System Biosciences). To generate a multicistronic tagging construct, the copGFP–polyA cassette was removed by EcoRI and NruI digestion, and the vector backbone was reconstituted via gBlock cloning to introduce a P2A sequence. A second gBlock encoding the mNeonGreen open reading frame followed by a WPRE element was inserted using an XhoI site and Gibson Assembly. For homology-directed repair (HDR)–mediated C-terminal tagging of CTIP2, the donor vector was sequentially digested with NheI and BamHI to allow insertion of the homology arms. The 5′ homology arm (420 bp) corresponded to the terminal exon of CTIP2, excluding the stop codon. The 3′ homology arm (420 bp) contained an in-frame P2A–mNeonGreen sequence, followed by the WPRE element and a loxP-flanked puromycin resistance cassette, resulting in the donor plasmid pTG-CTIP2:mNeonGreen. To generate CTIP2:mNeonGreen reporter lines in the fAD backgrounds, previously established APP isogenic hiPSC lines (AT, AV, SWE, and KO) were nucleofected with both pTG-Cr-CTIP2:mNeonGreen and pTG-CTIP2:mNeonGreen. Selection, expansion, and Cre-mediated excision of the selection cassette were performed as described above. Fluorescence-activated cell sorting (FACS) was performed 96 hours post-transfection to enrich for mNeonGreen-positive, mRuby-negative cells, thereby excluding false-positive fluorescence from transient pCAG-Cre:GFP expression. Clonal isolation and sequencing-based validation were carried out as previously described. Correct reporter localization was further confirmed by co-localization of endogenous mNeonGreen fluorescence with anti-CTIP2 immunostaining in dissociated cortical cultures. At least two independent positive clones per genotype were expanded for downstream experiments.

### Differentiation of hiPSCs into cortical spheres

hiPSCs were dissociated with Accutase (STEMCELL Technologies) for approximately 2 min at 37 °C and a total of 8 × 10^6^ hiPSCs were seeded into 10 cm^2^ ultra-low (ULA) attachment dishes (Corning) containing 10 ml mTeSR Plus medium supplemented with 10 µM Y-27632 (ROCK inhibitor; STEMCELL Technologies). Cultures were maintained at 55 rpm on an orbital shaker (VRW International) at 37 °C and 5% CO₂, as previously reported ^90^. Human iPSCs were maintained as undifferentiated pluripotent spheres in suspension with daily medium changes until reaching ∼50–100 µm in diameter. Cortical differentiation was initiated at day 0 (D0) by replacing 75% of the medium with differentiation medium consisting of 50% Advanced DMEM and 50% Neurobasal medium (Thermo Fisher Scientific), supplemented with 1% N2 (Life Technologies), 2% B27 without vitamin A (Invitrogen), 0.1 mM 2-mercaptoethanol (Life Technologies), 1% GlutaMAX (Thermo Fisher), and 200 µM ascorbic acid (Sigma-Aldrich). Medium was exchanged every other day by removing ∼50% of the volume after spheres settled by gravity (∼5 min). To promote dorsal forebrain specification, the differentiation medium was supplemented from D0 to D12 with the WNT inhibitor XAV939 (2 µM; Stemgent), the TGF-β/activin inhibitor SB431542 (10 µM; BioTechne, 1614/10), and the BMP inhibitor LDN-193189 (Hölzel, M1873). From D12 to D45, 20 ng/ml brain-derived neurotrophic factor (BDNF; Qkine, QK050) was added, and from D20 onward, 20 ng/ml glial cell line-derived neurotrophic factor (GDNF; Qkine, QK051) was included. Half of the medium was replaced every other day. For downstream experiments, cortical spheres were dissociated into single cells. Spheres were collected from spinner flasks and incubated in 2 ml of 0.25% trypsin supplemented with DNase I (50 µg/ml; Worthington Biochemical) for 7 min at 37 °C. Digestion was quenched with an equal volume of fetal bovine serum (FBS; Sigma-Aldrich) containing ovomucoid (1:5 dilution) and DNase I (0.5 mg/ml). Spheres were gently triturated five times using a 5-ml serological pipette, centrifuged at 800 rpm for 5 min, and resuspended in 5 ml of dissociation buffer composed of PBS supplemented with 5% FBS, 25 mM glucose, 5 mM GlutaMAX, papain (5 µg/ml; Worthington), and DNase I (14 µg/ml). Further mechanical dissociation was performed by gentle pipetting. Dissociated neurons were centrifuged (800 rpm, 5 min), resuspended in terminal differentiation medium containing BDNF (20 ng/ml), GDNF (20 ng/ml), and 1 µM cytosine β-D-arabinofuranoside (Ara-C), and passed through a 40-µm cell strainer (Corning). Cells were counted using an automated counter (Bio-Rad) and plated onto multiwell plates pre-coated overnight at 37 °C with poly-D-lysine (50 µg/ml; Sigma-Aldrich) in borate buffer (20×; Life Technologies), followed by a 4-hour incubation with fibronectin (50 µg/ml; Sigma-Aldrich) and laminin (4 µg/ml; BD Biosciences). Terminal differentiation medium was replaced every 3 days.

### Generation of cortical organoids (CxOs)

Cortical organoids were generated as previously described ^50^. Briefly, hiPSCs were dissociated using Accutase, and 6,000 cells per well were seeded into ULA attachment 96-well plates (Corning) containing 150 µl of mTeSR Plus medium (STEMCELL Technologies) supplemented with 50 µM Y-27632 (ROCK inhibitor). Plates were incubated at 37 °C with 5% CO₂ for 24 hours to allow spontaneous formation of embryoid bodies (EBs). For the first two days, EBs were fed by replacing 50% of the medium with fresh mTeSR Plus supplemented with 50 µM Y-27632 and fibroblast growth factor (FGF; 10 ng/ml; Life Technologies). The medium was then switched to mTeSR Plus without supplements, and daily half-medium changes were performed until EBs reached ∼600 µm in size. At this stage, the medium was replaced with cortical differentiation medium. After three days, when EBs exhibited smooth edges and a translucent outer surface - hallmarks of neuroectodermal differentiation - and regions of radially organized pseudostratified epithelium became visible, they were considered ready for Matrigel embedding. Embedding was performed as previously described in detail ^92^. Up to 48 embedded organoids were transferred to 10-ml ULA attachment culture dishes (Corning) and maintained on a horizontal shaker (VWR International) at 60 rpm. Medium was changed every two days, and organoids were cultured for up to 180 days.

### Immunocytochemistry and image analysis

hiPSCs and dissociated cortical cultures were fixed in 4% paraformaldehyde (PFA) for 20 min at room temperature, washed three times with PBS, and blocked for 1 h in a solution containing 10% normal goat serum (NGS) and 0.1% Triton X-100 in PBS. Cells were incubated overnight at 4 °C with primary antibodies diluted in blocking solution. After washing in PBS, samples were incubated with Alexa Fluor–conjugated secondary antibodies (Life Technologies) for 1 h at room temperature. Nuclei were counterstained with Hoechst 33342 (Life Technologies). Images were acquired using an automated Operetta CLS confocal high-content imaging system (PerkinElmer) with a 20× water objective. Image quantification was performed using Columbus Image Data Storage and Analysis software (PerkinElmer). A size- and morphology-based threshold defined by Hoechst staining was applied to exclude apoptotic nuclei from analysis. Neurons were identified based on MAP2 fluorescence, and following segmentation of nuclei and cell bodies, fluorescence intensities of the indicated antibody-labeled proteins were quantified as previously described ^41^. For immunostaining of CxOs, samples were fixed overnight at 4 °C in 4% PFA, washed in PBS, and cryoprotected in 15% sucrose overnight followed by 30% sucrose for an additional 24 h. Organoids were then embedded in Tissue-Tek OCT compound (Thermo Fisher Scientific) and snap-frozen. Cryosections (10 µm thickness) were prepared using a Leica CM1950 cryostat and collected on glass slides. Sections were rinsed in PBS, permeabilized, blocked, and immunostained using the same protocol as for dissociated cells. Images were acquired using either a Zeiss LSM 700 inverted confocal microscope or an Axio Scan Z1 slide scanner. The same imaging parameters were used across all CxOs derived from all hiPSC lines for each of the antibodies used. Maximum intensity projections of z-Stacks were made and stitched with 10% tile overlap by Hoechst33342 as reference channel with Global Optimizer option, using ZEN 3.2 blue (Zeiss). Automated time-lapse live-cell imaging was performed using an Operetta CLS high-content imaging system (PerkinElmer). A total of 80 × 10^4^ dissociated cortical neurons per line were seeded per well in 96-well plates and maintained in cortical differentiation medium (Thermo Fisher Scientific) supplemented with 10 ng/ml BDNF and GDNF. On days 3 and 14 after plating, cells were incubated with 10 nM calcein red (Invitrogen) for 30 min to label viable cells. Prior to imaging, cells were washed three times with cortical differentiation medium. Image analysis was conducted using Columbus Image Data Storage and Analysis software (PerkinElmer). Calcein-positive neurons were identified and quantified based on morphological parameters including cell body size, neurite length, cellular compactness, and fluorescence intensity. Primary antibodies used: CTIP2 (Abcam ab18465, rat), MAP2 (Lifespan Biosciences LS-C61805, chicken), OCT4 (Cell Signaling 2750 Rabbit), SOX2 (Invitrogen, 14-9811-82, rat), amyloid precursor protein (Abcam, ab32136, rabbit), PHF1 (Abcam ab184951 p-Tau Ser396/Ser404, rabbit), mouse β-amyloid (Biolegend, SIG-39320, 6E10), cathepsin B (Cell Signaling, 31718, rabbit), SDHB (Sigma-Aldrich, HPA002868, rabbit), TRXR2 (TXNRD2, sc-365714, mouse) rabbit β-tubulin (Abcam, ab6046), mouse β-actin (Cell Signaling Technology, 3700S). Congo red solution was used following manufacturer instructions (Sigma-Aldrich, HT603).

### Enzyme Linked Immunosorbent Assay (ELISA)

hiPSCs or differentiated neurons were seeded in 24-well plates, and cortical spheres were cultured in ultra–low attachment 6-well plates in their respective media. After three days, half of the culture medium from each well was collected, supplemented with protease inhibitors, and stored at −80 °C for subsequent analysis. Quantification of Aβ42 and Aβ40 was performed using a commercially available ELISA kit according to the manufacturer’s instructions (Life Technologies). To measure intracellular Aβ42 and Aβ40 levels, cells or cortical spheres from the corresponding wells were lysed in RIPA buffer (Sigma-Aldrich) supplemented with 2% SDS (Sigma-Aldrich) and protease inhibitors (Santa Cruz Biotechnology). Lysates were homogenized, heated at 95 °C for 10 min, and stored at −20 °C until use. Total protein concentration was determined using the BCA assay (Pierce) according to the manufacturer’s instructions. Aβ42 and Aβ40 peptide levels were normalized to total protein content, and values are reported as the amount of peptide per microgram of total protein.

### Flow analysis and FACS of CTIP2+ cortical neurons

To quantify the percentage of CTIP2:mNeonGreen⁺ neurons in each fAD line, dissociated day 45 cortical spheres were analyzed by fluorescence-activated cell sorting (FACS) using a BD FACSAria II+ flow cytometer. Approximately 2 million cells per sample were processed. Forward scatter (FSC) and side scatter (SSC) parameters were used to exclude debris and dead cells. Data were analyzed using FlowJo software. Each experiment included three technical replicates.

### Western Blot analysis

Cells were lysed in RIPA buffer (Thermo Fisher Scientific, 89900) supplemented with complete protease and phosphatase inhibitors, as described above. Protein extracts were separated on AnykD Criterion TGX Precast Midi Protein Gels (Bio-Rad, 5671124, 5671123) and run in 1× Tris-Glycine SDS Running Buffer (Thermo Fisher Scientific, LC26755) at 60 V for ∼30 min followed by 100–110 V for ∼2 h. Gels were equilibrated in 1× Tris/Glycine Transfer Buffer (Bio-Rad, 1610734) for ∼5 min prior to transfer. Proteins were transferred to membranes using the Trans-Blot Turbo Transfer System (Bio-Rad, 1704150) and pre-assembled Trans-Blot Turbo Transfer Packs (Bio-Rad, 1704157). Following transfer, membranes were rinsed with distilled water and stained with Ponceau to confirm uniform protein loading. Membranes were then blocked for 1 h at room temperature in 5% nonfat dried milk (PanReac Applichem ITW Reagents, A0830) prepared in 1× TBST (ChemCruz, sc362311) with gentle agitation. Primary antibodies were incubated overnight at 4 °C in blocking buffer. The next day, membranes were washed three times for 10 min each in 1× TBST, incubated with the appropriate HRP-conjugated secondary antibody diluted in 5% milk at room temperature, and washed again. Signal was developed using SupraSignal West PICO PLUS Chemiluminescent Substrate (Thermo Fisher Scientific, 34580), exposed to X-ray film (FUJI, 4741019289), and developed using a Cawomat 2000 IR X-ray film processor. Densitometric analysis was performed on scanned autoradiographs using Quantity One software (Bio-Rad).

### Proteomics, Cell lysis and protein digestion

Day 45 cortical spheres were lysed in 8 M urea, 150 mM NaCl, 0.5% NP-40 in 50 mM EPPS (pH 8.0), and sonicated. The homogenate was clarified by centrifugation at 21,000 × g for 5 min. Proteins were reduced with 5 mM tris(2-carboxyethyl)phosphine (TCEP) for 30 min at room temperature (RT) and alkylated with 10 mM chloroacetamide for 30 min at RT in the dark. Methanol–chloroform precipitation was performed prior to protease digestion. Precipitated protein pellets were resuspended in 100 µl of 100 mM EPPS buffer (pH 8.5) and digested at 37 °C for 2 h with LysC at a 200:1 protein-to-protease ratio. Trypsin was then added at a 100:1 protein-to-protease ratio and samples were incubated for an additional 6 h at 37 °C. Tandem mass tag (TMT) 10-plex labeling was performed using 100 µg of peptides per sample, following previously described protocols ^93^. Labeled peptides were pooled and fractionated using basic pH reversed-phase (BPRP) HPLC, as previously described ^94,95^. Mass spectrometry data were acquired using an Orbitrap Fusion Lumos mass spectrometer (Thermo Fisher Scientific) coupled to a Proxeon EASY-nLC 1000 liquid chromatography system (Thermo Fisher Scientific). Approximately 2 µg of each peptide fraction was loaded and separated using a 2.5 h gradient from 6% to 32% acetonitrile in 0.125% formic acid, at a flow rate of ∼500 nL/min. Proteome analysis used Multi-Notch MS3-based TMT quantification ^96^ to reduce ion interference compared to MS2 quantification ^97^. The scan sequence began with an MS1 spectrum (Orbitrap analysis; resolution 120,000 at 200 Th; mass range 350−1400 m/z; maximum injection time 50 ms; automatic gain control (AGC) target 5×105). For MS2 analysis, precursors (mass range 400-1400 m/z) were selected based on a top 10 scan method. MS2 analysis consisted of collision-induced dissociation (Quadrupole ion trap analysis; Turbo scan rate; AGC 1.0×104; isolation window 0.7 Th; normalized collision energy (NCE) 35; maximum injection time 35 ms). Monoisotopic peak assignment was used, and previously interrogated precursors were excluded using a dynamic window (180 s ±7 ppm). Following the acquisition of each MS2 spectrum, a synchronous-precursor-selection (SPS) - MS3 scan was collected on the top 10 most intense ions in the associated MS2 spectrum. MS3 precursors were fragmented by high-energy collision-induced dissociation (HCD) and analyzed using the Orbitrap (NCE 65; AGC 1.5×105; maximum injection time 150 ms, resolution was 50,000 at 200 Th). Raw mass spectra were processed using Proteome Discoverer (ThermoFisher – v2.3). Database searching was performed using SequestHT against the human Reference Proteome (UniProt database, downloaded in 05-2018), combined with a list of common laboratory contaminants, with a 15 ppm precursor mass tolerance and a 0.6 Da fragment mass tolerance. Peptide–spectrum matches (PSMs) were filtered using Percolator to achieve a false discovery rate (FDR) of 1%, and protein FDR validation was set to 1%. Reporter quantification filters included a co-isolation threshold of 40%, an average reporter S/N threshold of 10, and an SPS mass match percentage threshold of 70. Normalization mode was set to total peptide amount, and the maximum allowed fold change was set to 100 (default). Proteins quantified across both TMT sets (all 22 samples) were retained, and their quantification values were exported to Excel for further analysis. Supplemental Table 2 lists all quantified proteins along with their associated TMT reporter ion ratios used for quantitative analysis. Proteomics data were obtained from independent differentiation experiments. Statistical comparisons between conditions were performed in Prism or Perseus (v1.6.15) using unpaired two-tailed Welch’s t-tests (detailed parameters in Supplemental Table 2), or ANOVA (S0=0.585; 5% FDR correction). Raw data files are available upon request.

### scRNAseq / InDrop

Isogenic WT, AT, and SWE 45-day cortical spheres were dissociated as described above. Single-cell suspensions were encapsulated in droplets, and library preparation was performed according to a previously published protocol ^98,99^ with the following modifications to primer sequences to eliminate the need for custom sequencing primers: RT primers on hydrogel beads: 5’CGATTGATCAACGTAATACGACTCACTATAGGGTGTCGGGTGCAG[bc1,8nt]GTC TCGTGGGCTCGGAGATGTGTATAAGAGACAG[bc2,8nt]NNNNNNTTTTTTTTTTTTTTTTTTTV- 3’; R1-N6 primer sequence (step 151 in the library prep protocol in [2])- 5’TCGTCGGCAGCGTCAGATGTGTATAAGAGACAGNNNNNN-3’; PCR primer sequences (steps 157 and 160 in the library prep protocol in [2])- 5’-AATGATACGGCGACCACCGAGATCTACACXXXXXXXXTCGTCGGCAGCGTC-3’, where XXXXXX is an index sequence for multiplexing libraries. 5’-CAAGCAGAAGACGGCATACGAGATGGGTGTCGGGTGCAG-3’.

The workflow included the following steps: (1) encapsulation, (2) photocleavage of linkers, (3) reverse transcription, (4) second-strand synthesis, (5) in vitro transcription (IVT), (6) fragmentation, (7) second reverse transcription, (8) PCR amplification, and (9) library quality control. Libraries from each isogenic line were prepared in biological duplicate (n = 2 × 3 libraries), pooled based on molar concentration, and loaded at 2.07 pM on a NextSeq 75 (Illumina). Sequencing was performed with 26 bp for Read 1, 57 bp for Read 2, and 8 bp for Index 1. Data processing was performed using Cell Ranger (v1.2) (10x Genomics) for sample demultiplexing, barcode processing, and UMI counting. Reads were aligned to the GRCh38 reference genome (v12), and a digital gene expression matrix was generated for each experiment using default parameters ^100^. Basic processing and visualization of scRNA-seq data were conducted using the Seurat package (v2.3) in R (v3.3.4) ^101–103^.

### Clustering of Single-Cells

We analyzed cells of 3 samples: Wild Type (WT), A673V (AV) and Swedish Mutation (SWE). The filtered raw read counts were used as input for Seurat V3.1.4 (Stuart et al., 2019) by removing the following cells and genes; (i) cells with less than 200 unique transcripts (nFeature_RNA) and 500 total transcripts (nCount_RNA), (ii) genes detected in less than 5 cells, (iv) for AV and WT, we removed cells with more than 10000 nCount_RNA and 4000 nFeature RNA, for SW, we removed cells with more than 5000 nCount_RNA and 2000 nFeature_RNA. Following quality controls, 3643, 2858 and 6849 cells for WT, IT and SW datatsets, respectively, remained for further analyses. For each data, a Seurat object was generated by Normalizing (NormalizeData) and scaling data (ScaleData with vars.to.regress = c(“nCount_RNA”, “nFeature_RNA”) and calculating the first 20 PCs. Top 2000 variable genes were used to integrated datasets, followed by Normalization, Scaling, Clustering of the data. In total, 22 clusters were identified with resolution 1.

### Annotating Cell Types

**C**ell types were annotated in two steps: (i) identification of main cell types, and (ii) annotation of subtypes within each main cluster, when applicable. Following clustering and marker gene analysis, we identified five main cell types: VIM-, PTN-, FABP7-, and ID4-expressing cells as radial glia; RTN1-, DCX-, ELAVL3-, and SV2A-expressing cells as neurons; PCNA-, MKI67-, and TOP2A-expressing cells as proliferating cells; and TTR-expressing cells as choroid plexus. Four clusters (8, 19, 20, and 21) were annotated as low-quality cells due to enrichment for mitochondrial or ribosomal protein genes. Subtype annotation of each main cluster resulted in the final identification of 22 distinct cell clusters. We used the top 20 marker genes, along with known marker genes, to annotate neuronal subclusters. Subtypes were classified as follows: outer radial glia (oRG; HOPX, PEA15, LGALS3BP, MOXD1), apical radial glia (aRG; EMX1, SOX2, HES1), proliferating cells (Prolif; PCNA, MKI67, TOP2A), neuroblasts (NB; DCX, PCNA, ASCL1), immature neurons (immN; DCX), inhibitory neurons (inhN; GAD1, GAD2, SST, VIP), and excitatory neurons (ExN; SLC17A6, SLC17A7, NEUROD2, NEUROD6, FEZF2). (See Supplementary Figure 3.)

### Electrophysiological recordings and analysis

Before introducing the organoid, the MEA array was secured within the MEA Amplifier (MEA1060-Inv-BC, Multichannel Systems), which was mounted on the manual stage of an inverted Eclipse Ti-E/B microscope (Nikon) equipped with an interline CCD camera (Clara, Andor). Bright-field images at 2× magnification were taken to document electrode coverage by each organoid. The system was calibrated to 37 °C prior to recording. Cortical organoids were placed onto a 60-3DMEA250/12/100iR-Ti-gr multi-electrode array (Multichannel Systems). Organoids were transferred using a wide-bore 1250 µl pipette tip and positioned using a sterilized rat vibrissa. To enhance electrode–tissue contact without flattening the organoid, a modified slice grid (ALA-HSG-MEA-5b) was placed over the organoid, with the net elevated by 0.5 to 2 mm (adjusted in 0.5 mm increments depending on organoid diameter). Recordings were conducted once per week from week 6 to week 26 of organoid development using MC_Rack software (v4.6.2, Multichannel Systems). Electrophysiological signals were recorded for 10 minutes per organoid at a sampling rate of 40 kHz and a gain of 1100×. Only raw data were saved. During acquisition, signals were bandpass filtered (300–3000 Hz; Butterworth, 2nd order), and action potentials were detected using a threshold of –6 standard deviations with a 2 ms dead time. Detected spikes were visualized as overlays of 25 consecutive events from –2 to +5 ms relative to threshold crossing. Bursting activity was assessed through a continuous rate analysis using 500 ms bins. Live assessments of electrode activity were performed blind to genotype. Experimenters counted the number of electrodes covered by the organoid, as well as those exhibiting spiking or bursting activity during the recording session.

### Brain tissue immunostaining

Paraffin-embedded sections of the BA9 prefrontal cortex were obtained from the New York Brain Bank at Columbia University Irving Medical Center. Immunohistochemistry (IHC) was performed according to previously published protocols^104–107^. Sections were incubated with primary antibodies against SDHB (Sigma-Aldrich, HPA002868, rabbit polyclonal, 1:500) and TRXR2 (TXNRD2, Santa Cruz, sc-365714, mouse monoclonal, 1:500), followed by incubation with appropriate Alexa Fluor-conjugated secondary antibodies (Thermo Fisher, 1:500). Nuclei were counterstained with Hoechst. Confocal images were acquired using a Zeiss LSM800 microscope with fixed acquisition parameters. When applicable, Z-stacks were projected as maximum-intensity images for analysis. Image acquisition and quantification were performed blinded to experimental conditions. Data distributions were assessed using Shapiro–Wilk, D’Agostino–Pearson, Anderson–Darling, and Kolmogorov–Smirnov tests. Statistical comparisons were performed using the non-parametric Mann–Whitney U test. Quantitative data are reported as mean ± standard error of the mean (SEM).

### Proteomics of AMP-AD/ROSMAP

Proteomics data from generated in autopsied brains from the ROS/MAP cohort were downloaded from the AD knowledge portal (https://www.synapse.org/Synapse:syn17015098). Data generation, pre-processing and protein quantification have been described previously^108–111^. After regressing out the effect of age and sex from protein levels, we used TAMPOR (Tool for Analyzing and Mitigating Proteomics Batch Effects and Outliers Robustly) to correct batch effects in the proteomics dataset. TAMPOR was configured to use all non-GIS channels for correction, with median-based central tendency adjustment and 250 iterations for robust data normalization. Processing was parallelized across eight threads, with a minimum batch size of five samples to prevent overfitting. We then tested the association of AD neuropathological features with protein levels using linear and logistic regression models adjusting for age, sex and APOE-ε4 genotypes.

### Statistical Analysis

Statistical analyses were performed using GraphPad Prism version 10.3.1 (GraphPad Software, Inc.). The normality of residuals was assessed using the Shapiro–Wilk test. For comparisons involving a single variable and two groups (e.g., genotype), a paired two-tailed t-test was used when data were normally distributed. If residuals did not follow a Gaussian distribution, the nonparametric Wilcoxon matched-pairs signed-rank test was applied. For comparisons across three or more groups involving a single independent variable, one-way ANOVA was conducted, followed by Dunnett’s multiple comparisons post hoc test. For experiments involving multiple groups and more than one independent variable, two-way ANOVA was used, followed by multiple comparison post hoc tests as indicated in the corresponding figure legends. A 95% confidence interval was used for all statistical tests. All experiments represent the results of at least three independent biological replicates. Graphs depict mean ± standard error of the mean (SEM), unless otherwise noted. Statistical significance was defined as follows: * p < 0.05, ** p < 0.01, *** p < 0.005, **** p < 0.001, ***** p < 0.0001. “ns” indicates not significant.

## Acknowledgments.

We thank Prof. Christian Haass for insightful feedback on the manuscript. We thank the Single Cell Core at Harvard Medical School (Boston, MA) for support with single-cell RNA-seq library preparation and Lena Hersemann (MPI-CBG Scientific Computing Facility) for support with mass spec data analysis. We thank Dr. Andrew F. Teich, Delaney Flaherty and New York Brain Bank for post-mortem human brain tissue, Taub Institute Imaging Platform, the contributors, who collected samples used in this study, and the patients and families for their participation. We thank Silke White (DZNE Imaging platform) for her support and training on multiple microscopes and Beate Knauer (CRTD Electrophysiology Facility, TU Dresden) for providing infrastructure and assistance on performing MEA-recordings.

## Author Contributions

TG: Conceptualization, data generation and curation, formal analysis, methodology, writing. MIC: scRNAseq analysis. AO: proteomic data acquisition and analysis. AC: methodology. FDP: scRNAseq data acquisition. SK: electrophysiology data acquisition and analysis. MNQ: ROSMAP proteomics analysis. BV: ROSMAP proteomics analysis. JP: proteomics methodology. MM: conceptualization, editing. LLR: funding acquisition, supervision. CK: formal analysis, supervision, funding acquisition, writing. NRM: conceptualization, data generation and curation, formal analysis, supervision, writing, funding acquisition.

