## Supplementary figures and images for "Isogenic cortical organoids enable precision targeting of APP variant-specific pathways in Alzheimer’s disease"

### Supplemental figure 1

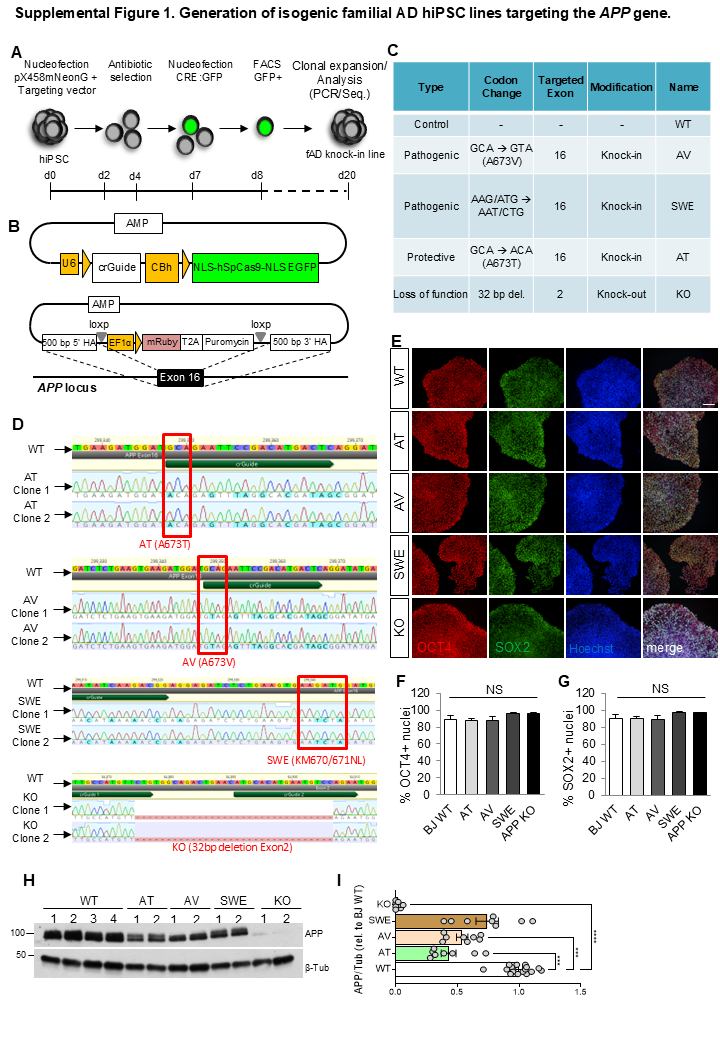

### Supplemental figure 2

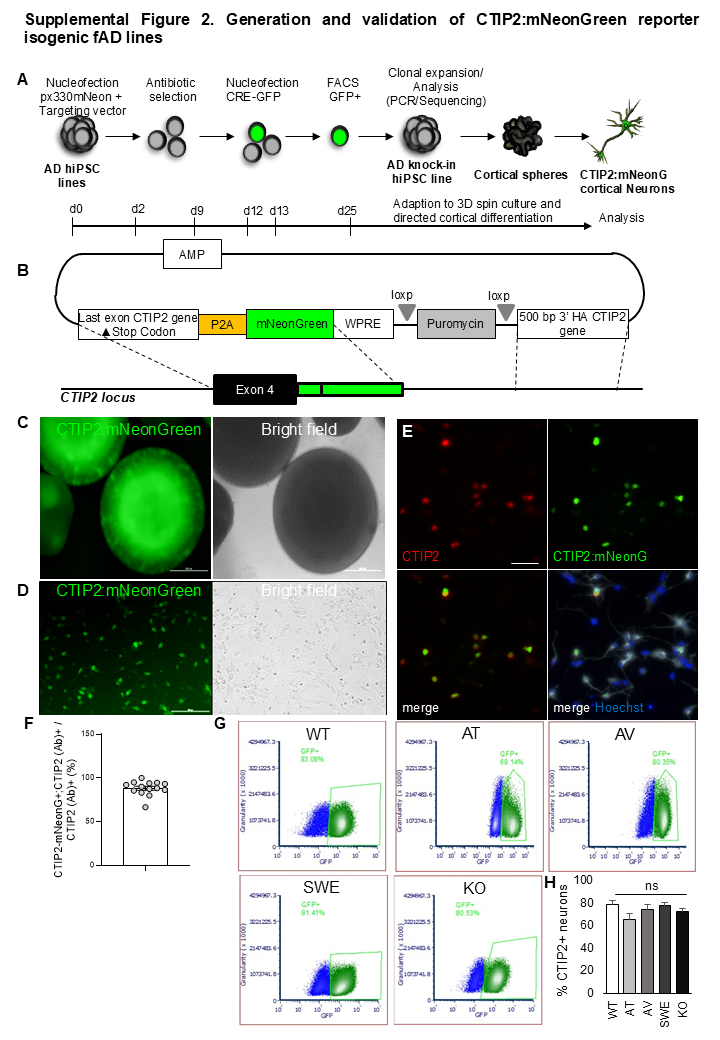

### Supplemental figure 3

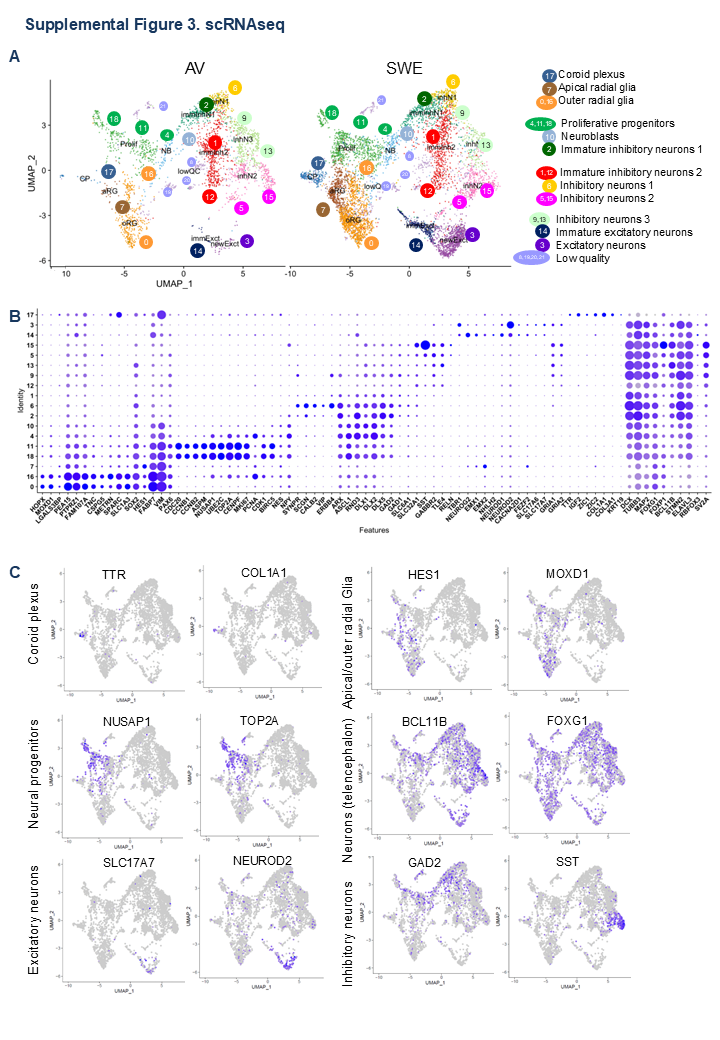

### Supplemental figure 4

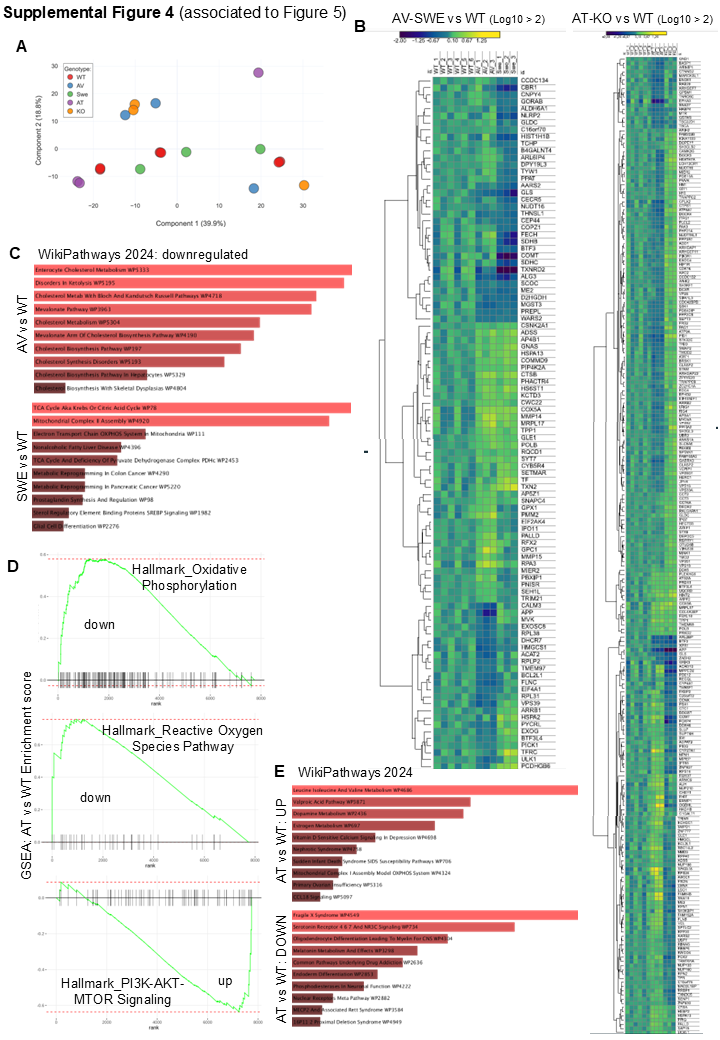

### Supplemental figure 5

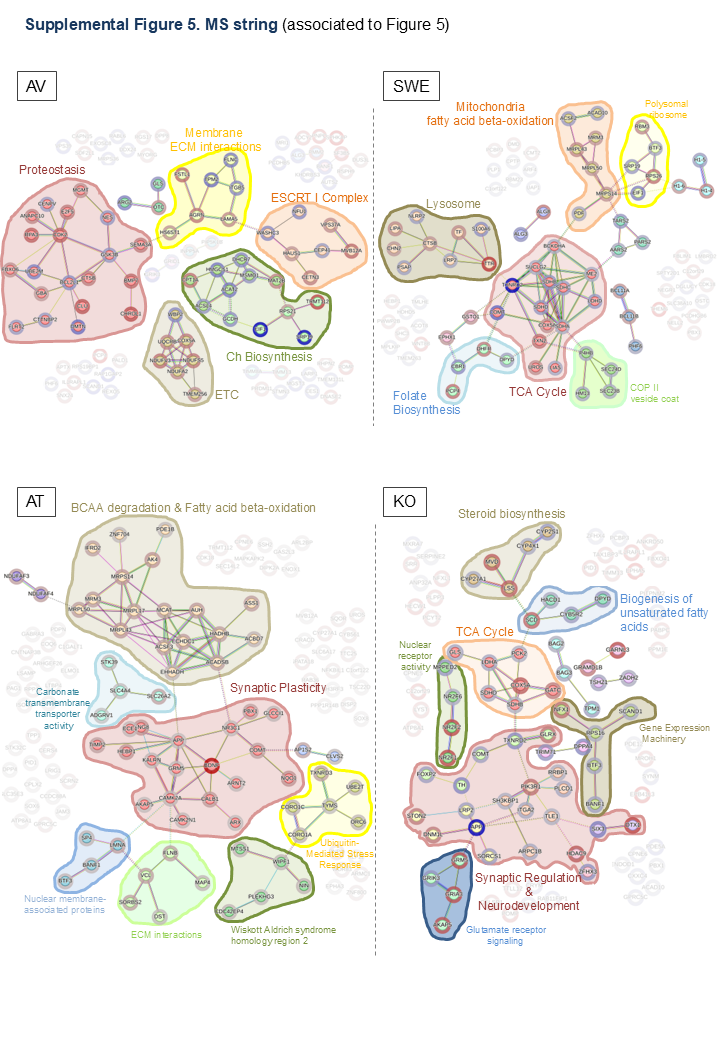

### Supplemental table

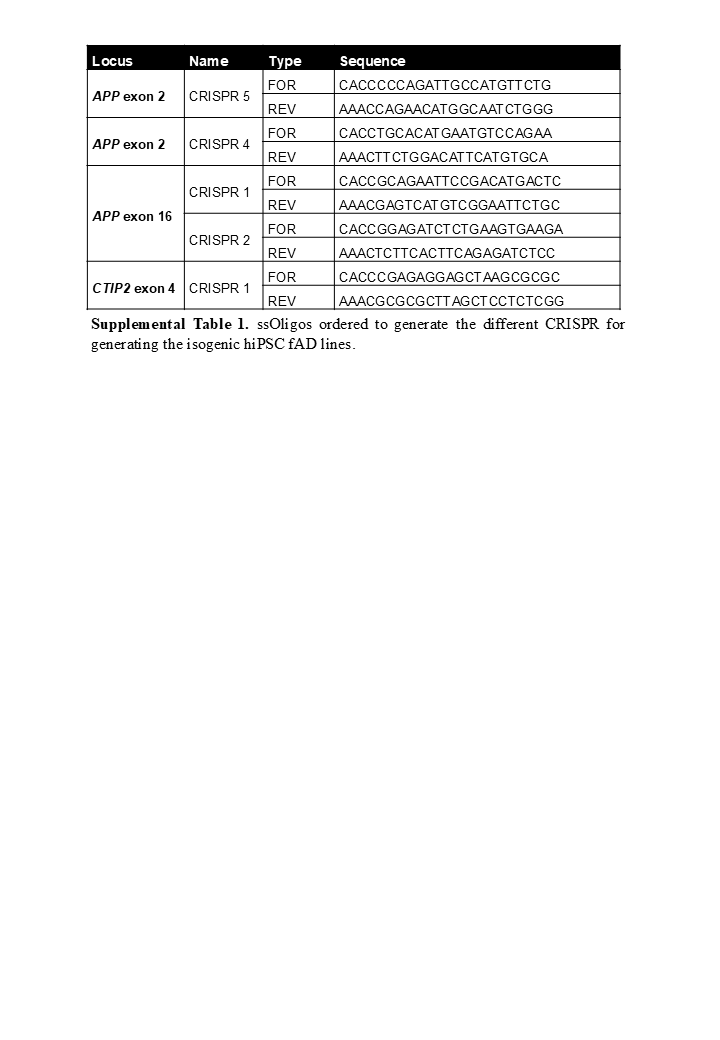
